# Improved HaloTag for analyses of translocation of type III secretion system effector proteins

**DOI:** 10.64898/2026.05.31.729057

**Authors:** Verena Nadin Fritsch, Michael Holtmannspötter, Michael Hensel

## Abstract

Effector translocation during host-pathogen interactions is a prerequisite for the entry of *Salmonella* into non-phagocytic cells and establishment of a replication permissive intracellular niche. Deciphering the dynamics and kinetics of translocation and subcellular localization demands live-cell imaging and tagging approaches that do not introduce detection delays or perturb the translocation process via the type III secretion system (T3SS). Effector fusions with self-labelling enzymes (SLE), such as HaloTag, allow localization and tracking at high temporal and spatial resolution. However, interference with T3SS-dependent translocation has hampered analyses of the process of translocation and early subcellular distribution and dynamics. Herein, we report that amino acid substitutions of the HaloTag can reduce the thermodynamic stability, resulting in less steric hindrance during translocation of effector-HaloTag fusions by the T3SS in mammalian cells. The top variant, HT-SP5, showed reduced retention in *Salmonella*, enabling more sensitive and earlier detection of translocated effector proteins of the SPI1 and SPI2 T3SS of *Salmonella* and of the T3SS effector Map of enteropathogenic *Escherichia coli* (EPEC). We applied the improved HaloTag HT-SP5 to single molecule tracking, and to follow effector protein dynamics in living host cells early after translocation by invading and intracellular bacteria. Taken together, the improved HaloTag variant HT-SP5 represents a robust and versatile SLE tag for dynamic real-time analyses of delivery and fate of T3SS-translocated effector proteins in living cells host. Application of HT-SP5 will facilitate research on effectors throughout the entire infection process at native effector levels to understand host-pathogen interactions.

## Introduction

Bacterial secretion systems mediate the export of diverse substrates, including toxins, adhesions and enzymes, that are crucial for bacterial survival and shaping of host-pathogen interactions (Jamali et al., 2025). Among secretion systems in Gram-negative bacteria, especially the injectisome-like secretion systems have gained special attention as promising target to inhibit bacterial virulence and pathogenesis (Blasey et al., 2023). In contrast to other secretion systems, these multi-subunit molecular nanomachines mediate the direct one-step translocation of effector proteins from the bacterial cytoplasm into the host cell cytoplasm. Once inside the cell, these effectors subvert cellular signalling pathways to promote immune evasion and intracellular bacterial survival. The pathogenesis of *Salmonella enterica*, the leading cause of bacterial foodborne illness globally (Wang et al., 2025), is for example intricately linked to the functionality of its two distinct type III secretion system (T3SS) (Mishra et al., 2025). While the T3SS encoded by genes in *Salmonella* Pathogenicity Island (SPI) 1 is required for invasion of non-phagocytic cells by orchestrating a series of cytoskeletal manipulations, the SPI2-encoded T3SS is essential for the formation of a replication-permissive niche within host cells and prolonged intracellular survival. SPI2-T3SS effectors mediate massive endosomal remodelling in host cells, resulting in formation of *Salmonella*-containing vacuoles (SCV) and *Salmonella*-induced filaments (SIF) (reviewed in Liss & Hensel, 2015).

Despite their structural conservation among Gram-negative bacteria (Cabezón et al., 2023; Carsten et al., 2022; Keck et al., 2024), the mechanism of T3SS translocation and especially subsequent fate of translocated effector proteins in host cells and interactions with host cell membranes remain insufficiently resolved. Use of fluorescent proteins such as GFP for elucidation of translocated is hampered by the narrow inner needle diameter of 15-35 Å (1.5-3 nm), which requires effectors to unfolded during translocation and refold within the host (Cabezón et al., 2023; Cornelis, 2006; Guzmán-Herrador et al., 2023; Wagner et al., 2018; Worrall et al., 2023). Especially the ease of unfolding under force (mechanical stability) and the thermodynamic stability, rather than the protein size per se, is the determining factor for effector translocation (Akeda & Galan, 2005; DaPron et al., 2026; LeBlanc et al., 2021; Radics et al., 2014). Consequently, the labelling of effector proteins for live-cell imaging to follow their translocation, subcellular localization and dynamics in the host proved challenging. Conventional fluorescent proteins, such as GFP, have been shown to be incompatible with these secretion machineries. Over the last years a few alternative imaging approaches have been described (reviewed in Fritsch & Hensel, 2025). Of these, self-labelling enzymes (SLE) have proven well-suited to trace effector proteins on a molecular level in living cells.

SLE tags, such as SNAP-Tag and HaloTag, are an innovative labelling technology with several advantages over fluorescent proteins, including higher brightness, greater photostability, and a broader spectrum of fluorophores, making them compatible with advanced fluorescence imaging techniques. Recently, we have demonstrated their feasibility for analyses of effector translocation into living cells and the dynamics of SPI2-T3SS effector proteins SifA, SseF and PipB2 in interaction with host cell endosomal membranes such as SCV and SIF. However, effector-HaloTag fusions showed limited translocation efficacy, particularly for SPI1 effectors. This resulted in incomplete restoration of virulence in the respective deletion strain, and low signals for the translocated portion (Göser et al., 2019), restricting our analyses to abundant effector proteins, and effector protein dynamics at later stages of infection. While smaller SNAP-Tag and CLIP-Tag fusions proved suitable for SPI1-T3SS effectors, the usability of HaloTag would allow multiplex analyses of effectors due to the orthogonality of these tagging strategies. Additionally, HaloTag remains the preferred tool due to its superior labelling kinetics (∼ 100-fold faster than SNAP-Tag) (Hoelzel & Zhang, 2020), enhanced brightness (∼2- to 6-fold higher labelling intensity than SNAP-Tag) and photostability, making HaloTag superior for super-resolution imaging (Erdmann et al., 2019). To overcome these constraints and enable the study of early translocation events and protein dynamics under native conditions, a more compatible labelling strategy is required. Motivated by recent achievement in fluorescence intensity, photostability and labelling kinetics of HaloTag (Frei et al., 2022; Ling et al., 2025; Miro-Vinyals et al., 2021), we set out to identify HaloTag alleles with improved compatibility in T3SS-mediated translocation. Here we report that HaloTag allele HT-SP5 with amino acid exchanges M175Y V245A L271D, is optimized for analyses of T3SS effector protein translocation. SPI1- and SPI2 effector-HT-SP5 fusions are efficiently translocated into host cells while retention in bacteria was nearly absent. HT-SP5 provides a powerful tool for analyses of T3SS effector translocation and dynamics in living host cells.

## Results

### Rationale for generation of improved HaloTag as translocation reporter

We demonstrated the use of HaloTag and other SLE as reporters for translocation and intracellular fate of T3SS effector proteins (Göser & Hensel, 2021; Göser et al., 2019; Göser et al., 2023). In these previous experiments with native HaloTag (i.e. HaloTag7), the intense labelling by HTL of bacterial cells and rather low signal of translocated effectors in host cells was observed (**Figure 1**). Only partial complementation of effector deletion mutant strains by episomal effector-HaloTag fusions was observed (Göser et al., 2019), indicating inefficient T3SS-dependent translocation of effector proteins fused to native HaloTag. Since previous studies demonstrated that targeted mutagenesis of the HaloTag resulted in altered characteristics, we hypothesized that certain mutant alleles of HaloTag affect the thermodynamic stability and (un)folding kinetics, leading to improved effector translocation. We created a set of fusions between candidate effector proteins and mutant alleles of HaloTag. As before (Göser et al., 2019; Göser et al., 2023), HaloTag was fused C-terminal to candidate effector in order to maintain N-terminal chaperone-binding domains and secretion signals. Focussing on previously reported HaloTag alleles with enhanced labelling kinetics and brightness (Kang et al., 2017; Miro-Vinyals *et al.,* 2021; Lampkin et al., 2024), we engineered six HaloTag variants featuring one to four amino acid (aa) exchanges. Furthermore, a short, universal GSGSGS sequence linked effector and HaloTag, and the HA tag was fused C-terminal to enable quantification of levels of effector-HaloTag protein independent from HTL labelling. Microscopic analyses and functional tests revealed no obvious performance alterations due to the linker exchange (**Figure 1A**, **Figure 2**).

**Figure 1:**
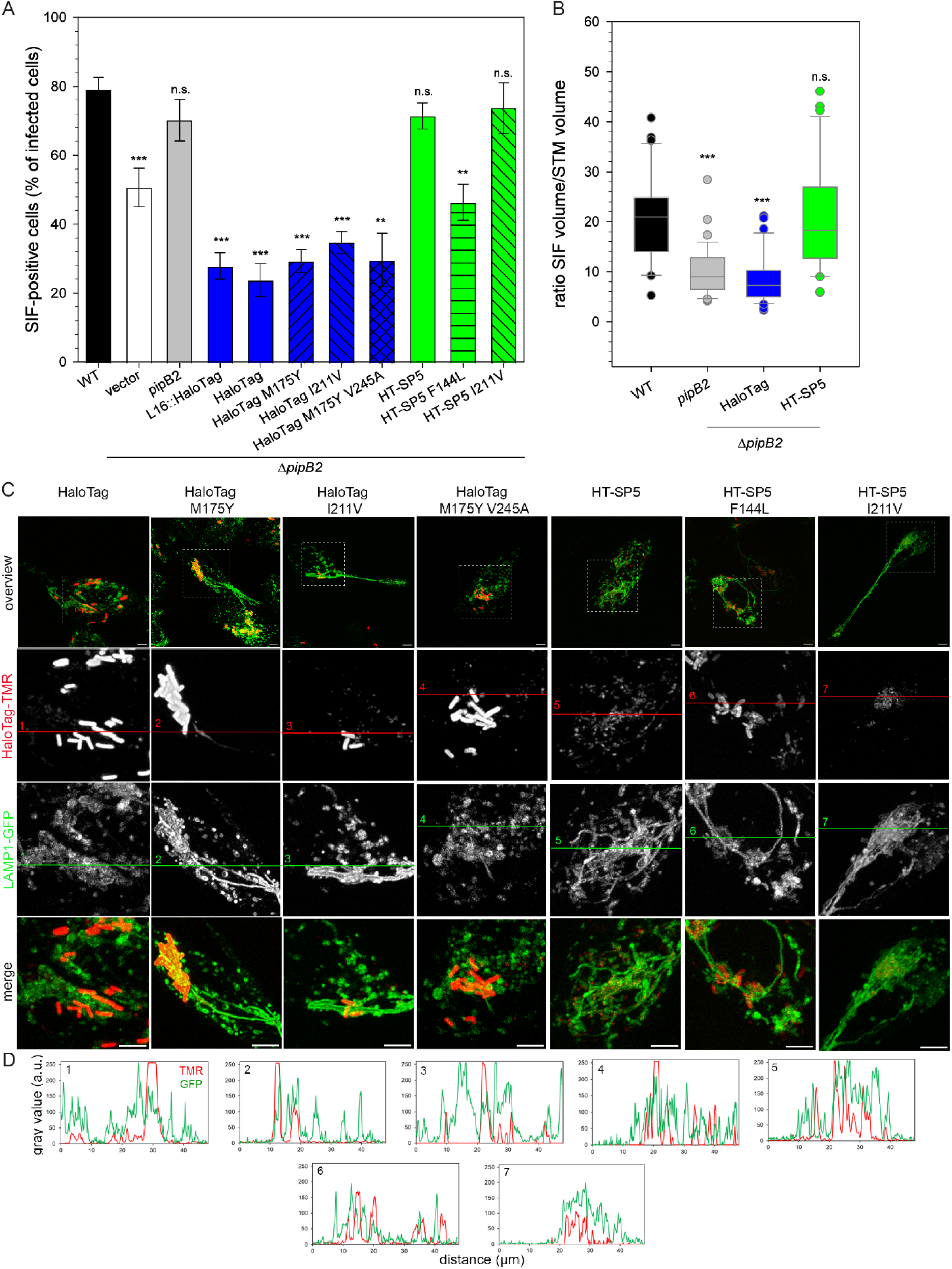
Performance of various HaloTag alleles in translocation of SPI2-T3SS effector proteins. HeLa LAMP1-GFP cells were infected with STM strains WT or Δ*pipB2*, each expressing sfmTurquoise2ox, complemented with plasmids for expression of *pipB2* fused to various HaloTag alleles. **A**) The percentage of SIF-positive cells was determined by LCI at 8 h p.i. of at least 100 infected cells in three biological replicates each. The means and standard deviations are shown. **B**) After three-dimensional reconstruction of infected cells by Imaris, the SIF network was quantified by determining the volume of interconnected LAMP1-GFP-positive tubules in relation to the volume of enclosed bacteria. Images representative for at least 30 analysed cells per condition are shown in **Figure S1**. Statistical significance determined by unpaired two-sided *t*-test is indicated as: n.s., > 0.05; *; p < 0.05; **, p < 0.01; ***, p < 0.001. **C**) HeLa cells stably expressing LAMP1-GFP were seeded in 8-well chamber slides. The next day, cells were infected at MOI 10 with STM Δ*pipB2* harbouring plasmids for expression of *pipB2* fused to various HaloTag alleles as indicated. LCI was performed at 8 h p.i. Labelling reactions using 1 µM HTL-TMR were performed directly before imaging for 30 min at 37 °C. Representative STM-infected host cells exhibiting SIF formation were selected for LCI. Scale bars, 10 µm. D) Line scans reveal signal intensity of translocated and intra-bacterial effector proteins (red) in relation to LAMP1-GFP (green).

**Figure 2:**
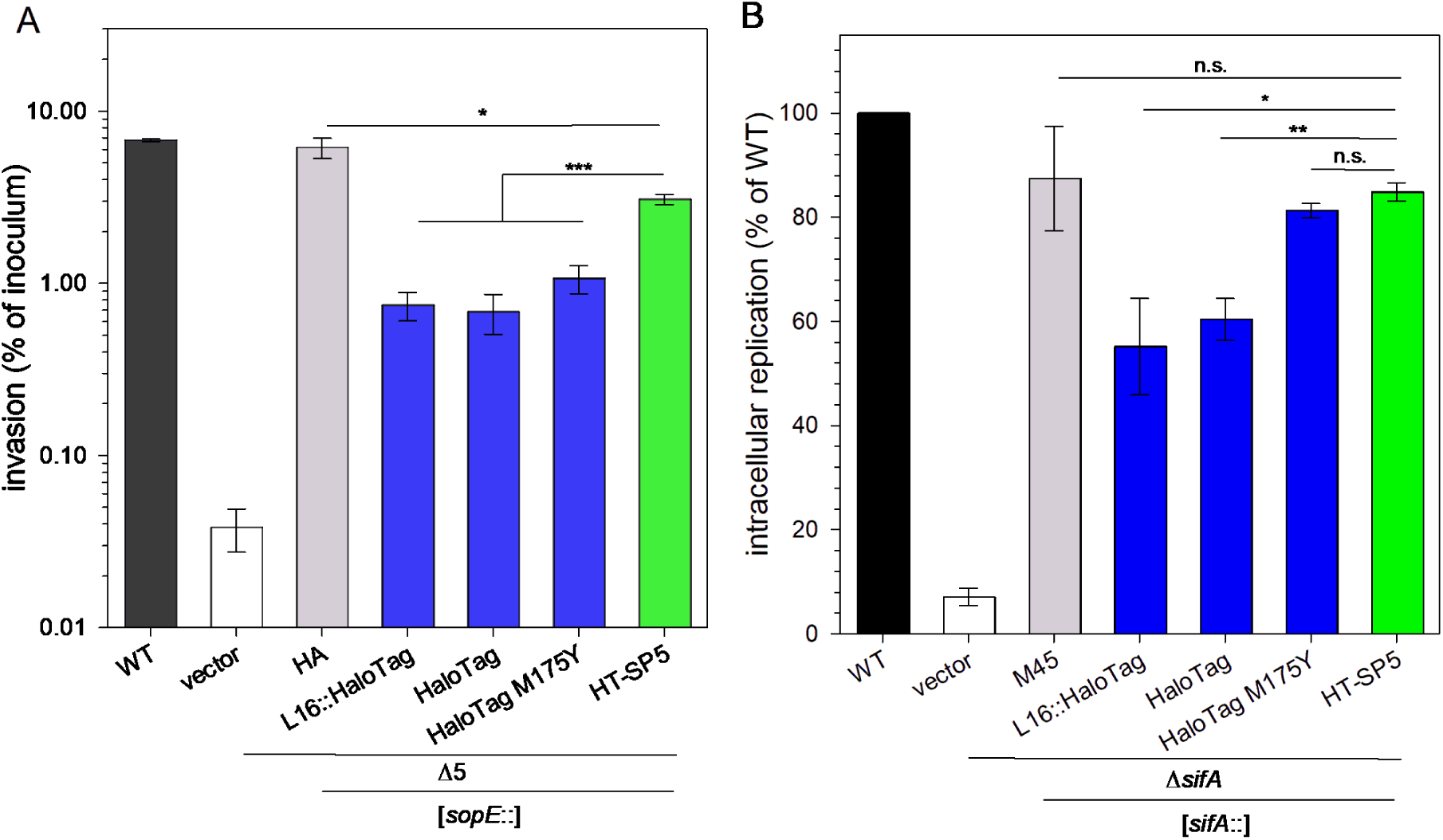
T3SS effector fusion to HaloTag allele HT-SP5 improves complementation of mutant strains deficient in invasion or intracellular replication. Invasion of HeLa cells (**A**), and intracellular replication in RAW264.7 macrophages (**B**) was determined by gentamicin protection assays. Functional complementation of mutant strains defective in invasion (Δ5 strain, deletion of SPI1-T3SS effectors *sipA*, *sopB*, *sopD*, *sopE*, and *sopE2*) or intracellular proliferation (Δ*sifA*), by plasmids for expression of HaloTag fusions to SPI1-T3SS effector *sopE* (A) or SPI2-T3SS effector *sifA* (B) were analysed after infection at MOI 5 (**A**) or MOI 10 (**B**). Medium containing 100 μg/ml gentamicin was used for 1 h to kill extracellular bacteria. Intracellular CFU counts were determined 1 h, 2 h and 16 h p.i., and invasion is the ratio of CFU recovered at 1 h and CFU in inoculum, while replication is the ratio of CFU recovered 16 h and 2 h p.i. Depicted are means and standard deviations of biological triplicates, with three technical triplicates each, and statistical significance was determined by unpaired, two-sided *t*-test as: n.s., > 0.05; *, < p0.05; **, p < 0.01; ***, p < 0.001.

### Selection of HaloTag variants with improved translocation efficiency

To evaluate HaloTag alleles regarding effector translocation and functionality, we quantified the proportion of infected cells showing formation of *Salmonella*-induced filaments (SIF) as proxy for SPI2-T3SS-dependent endosomal remodelling by STM (**Figure 1A**, **Figure S1**). SIF biogenesis is dependent on effector protein SifA, with contributions by SseF, SseG, PipB2, SseJ, and SopD2, and the deletion of *pipB2* resulted in 1.6-fold reduction of SIF-positive cells. Importantly, we noticed that complementation of the Δ*pipB2* deletion mutant with *pipB2*::HaloTag resulted in decrease of SIF-positive cells to 30% compared to 51% and 80% observed after infection with the deletion mutant and WT, respectively (**Figure 1A**). These results indicate, in accordance with our previous observations (Göser & Hensel, 2021; Göser et al., 2019) that effector-HaloTag fusions partially interfere with T3SS-dependent delivery, resulting in a partially block of T3SS and thereby inhibition of effector translocation. Strikingly, effector-fusions to two HaloTag variants, HaloTag M175Y V245A L271D (hereafter referred to as ‘HT-SP5’) and HaloTag M175Y V245A L271D I211V, produced similar phenotypes to the WT and HA-epitope-tagged version, which is known to interfere minimally with translocation by T3SS. Since PipB2 is known to be essential for the centrifugal extension of the SIF network, we quantified the SIF volume in dependence on the bacterial load for cells infected with STM strains WT, Δ*pipB2*, or Δ*pipB2* complemented with *pipB2*::HaloTag or *pipB2::*HT-SP5 fusions (**Figure 1B**). In accordance with previous reports (Knodler & Steele-Mortimer, 2005; Rajashekar et al., 2014), the SIF network appeared bulky with short SIF filaments in cells infected by STM Δ*pipB2*. In contrast, SIF network in cells infected by STM WT, or Δ*pipB2* expressing *pipB2*::HT-SP5 was elongated and extended distal from SCV to cell periphery, resulting in 2.5-fold higher SIF/STM volume ratios than observed for STM Δ*pipB2* complemented with *pipB2*::HaloTag (**Figure 1B**).

We next evaluated the performance of the effector-HaloTag fusions in live-cell imaging (LCI). HeLa cells infected by STM expressing effector-HaloTag fusions were treated with 1 µM HTL-TMR and imaged 8 h p.i. (**Figure 1CD**). In accordance with partial complementation of mutant phenotypes (**Figure 1A**), we observed strong signal accumulation within STM and weak signals of translocated effector proteins fused to HaloTag, HaloTag M175Y, HaloTag V245A, HaloTag M175Y V245A and HaloTag I211V. HaloTag allele M175Y V245A L271D F144L showed robust labelling of translocated effector proteins but still retained, unlike the HT-SP5, considerable amounts of non-translocated effector-fusions inside the bacterial cell. By contrast, only a very weak signal could be detected for *pipB2*::HaloTag M175Y V245A L271D I211V-expressing bacteria. Based on these results, we focussed subsequent analyses on most promising HaloTag allele HT-SP5.

### Enhanced translocation of functional SPI1 and SPI2 effector-HaloTag-SP5 fusions

Following on the finding of enhanced translocation of PipB2 fused to HT-SP5, we determined if fusions with other effector proteins would also result in more efficient delivery of these virulence factors than with the original HaloTag. Complementation of STM Δ*sifA* with plasmids for expression of *sifA*::HT-SP5 restored intracellular proliferation in macrophages to the same extent as observed for *sifA* with M45-tag only (**Figure 2B**). In contrast, intracellular replication of STM Δ*sifA* expressing *sifA*::HaloTag was reduced by almost 30%. LCI of further SPI2-T3SS effector proteins fused to HaloTag or HT-SP5 after labelling with HTL-TMR in living host cells (**Figure S2**) revealed strong intra-bacterial accumulation of SifA-HaloTag and SseJ-HaloTag. Conversely, SifA-HT-SP5 and SseJ-HT-SP5 fusions were efficiently translocated into the host cells and preferentially localized to SCV and SIF membranes. Importantly, we detected SifA-HT-SP5 decorated host endosomal vesicles near the growing SIF network. Although SifA is required for the remodelling of the host endosomal system, low translocation levels of SifA-HaloTag hampered detection of SifA decorated vesicles in earlier studies (Göser et al., 2023). To quantify translocation efficiency and retention, we determined the intra-bacterial effector-bound TMR intensity in a mutant deficient in SPI2-T3SS translocation (Δ*ssaV),* as well as total effector-bound TMR intensity in infected RAW macrophages and of the isolated bacterial fraction, respectively. We observed no statistic difference when measuring the fraction containing cells and intracellular bacteria (**Figure S4A**), or when performing LCI of Δ*ssaV* mutant expressing *pipB2*-HT-SP5 and *pipB2*-HT-SP5 (**Figure 4AB**), indicating similar expression and labelling levels of PipB2-HaloTag and PipB2-HT-SP5 8 h p.i. The isolated bacteria expressing *pipB2*-HT-SP5 showed a 2.3-fold lower TMR intensity (**Figure S4A**), confirming more efficient effector delivery into the host cell cytosol as observed in our LCI analyses.

Effector proteins of the SPI1-T3SS fused to HaloTag were translocated inefficiently and use of SNAP-tag was recommended (Göser et al., 2019). To determine whether the HaloTag allele HT-SP5 may improve performance, we analysed secretion dynamics of SopB fused to HaloTag or HT-SP5 (**Figure S3**). While a peak in SopB-HT-SP5 secretion was already detected after 4 h sub-culture, SopB fused to the original HaloTag appeared not until 6 h and reached its maximum after 8 h. Accordingly, high retention of SopB-HaloTag in bacterial cells was determined, which was absent for the SopB-HT-SP5 fusion protein, unless expressed in secretion-deficient STM Δ*invC*. Flow cytometry analyses of secretion competent STM further confirm the efficient and fast secretion of the labelled effector-HT-SP5 fraction, and intra-bacterial retention of effector-HaloTag fusions (**Figure S4BCD**).

To investigate *in vivo* functionally of SPI1-T3SS effector-HaloTag fusion proteins, we analysed the invasion of epithelial cells by STM Δ5 mutant strain deficient in SPI1-T3SS effector proteins SipA, SopA, SopB, SopE, and SopE2 without and with complementation by *sopE*::HaloTag fusions **(Figure 2A**). Although expression of the *sopE*::HT-SP5 fusion could not restore the attenuated invasiveness of the Δ5 strain to the level of the WT, the invasion was 4-fold elevated compared to the strain expressing the *sopE*::HaloTag fusion.

We further tested the translocation and subcellular localization of SipA, another representative effector protein of the SPI1-T3SS, upon fusion with HT-SP5. SipA fusion to initial HaloTag resulted in substantial amounts of effector protein retained inside the bacteria and only a weak signal of translocated effector protein could be observed at the poles of bacteria (**Figure 3**). In contrast, bacteria expressing *sipA*::HT-SP5 fusions showed low intra-bacterial signal intensities. Instead, translocated SipA was visible adjacent to the bacteria.

**Figure 3:**
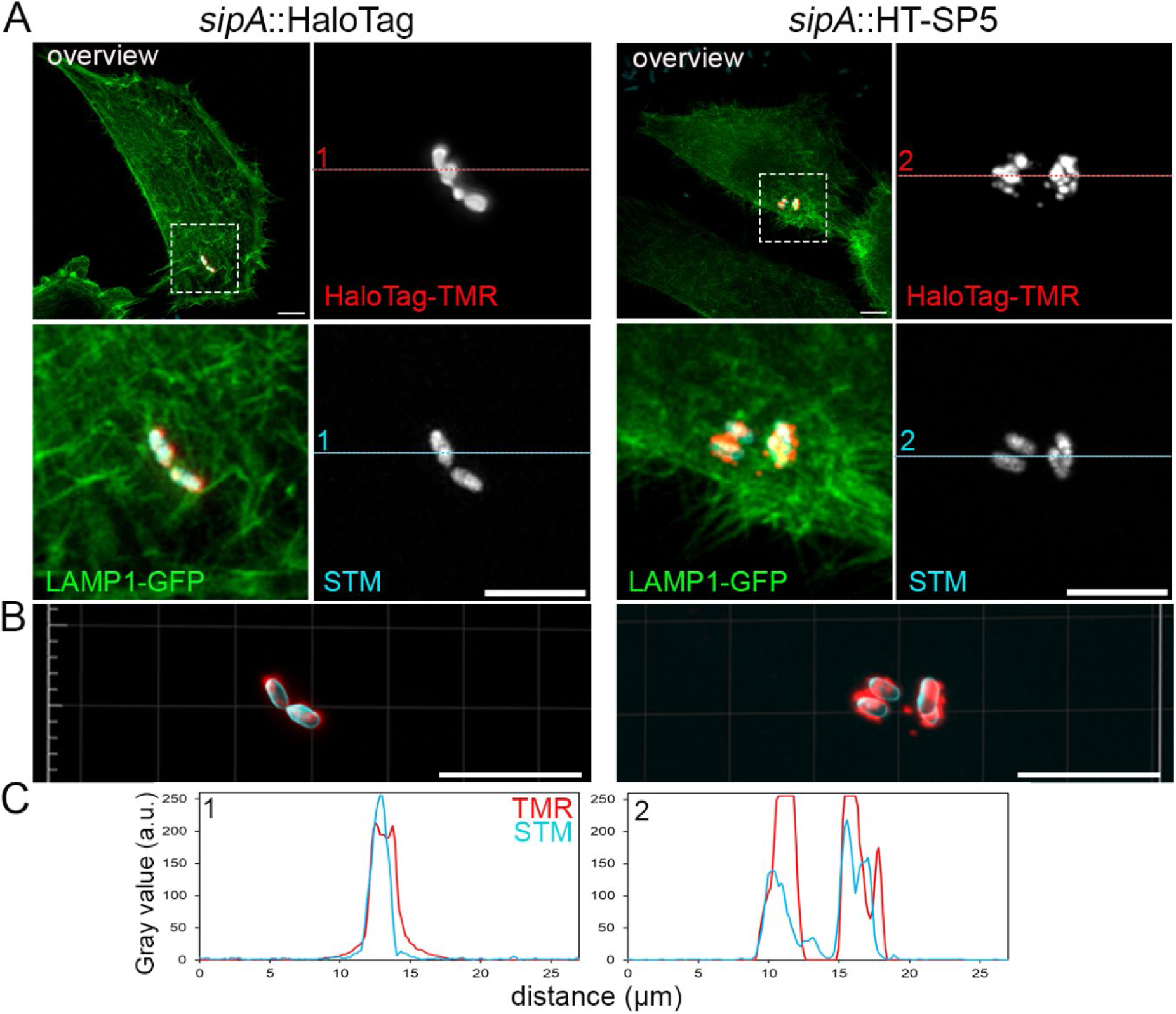
HaloTag allele HT-SP5 improves translocation and subcellular distribution of the SPI1-T3SS effector protein SipA. **A)** HeLa cells stably expressing LifeAct-GFP (green) were seeded in 8-well chamber slides. WT STM (light blue) expressing *sipA*::HaloTag or *sipA*::HT-SP5 were sub-cultured in LB for 3.5 h to induce SPI1-T3SS expression. During infection, 1 μM HTL-TMR (red) was added for 30 min, before cells were fixed by PFA and subjected to imaging using CLSM (Olympus LSM FV3000). **B)** 3D reconstructions of volumes of STM and translocated SipA-HaloTag fusions labelled by HTL-TMR in representative cells. Scale bars, 10 μm. **C)** Line scans for quantification of signal intensity of translocated and intra-bacterial effector proteins (red) in relation to STM (blue) for representative infected cells.

From these data, we conclude that HT-SP5 is highly compatible with T3SS-dependent translocation, allowing improved detection and analyses of functional effector proteins in living host cells.

### HT-SP5 exhibits lower labelling efficiency and decreased thermal stability

Increased signal intensities for translocated effector proteins may be caused by higher compatibility of HaloTag alleles with translocation by T3SS, enhanced efficiency of self-labelling with HTL, and/or increased signal intensity of the fluorophore of the HTL. The improved complementation of mutant strains by effector fusions to HT-SP5 (**Figure 1**, **Figure 2**) supports the first mechanism.

To evaluate the labelling efficiency and performance of HaloTag alleles independent from T3SS translocation, we measured TMR/GFP ratios in transfected HeLa cells co-expressing cytosolic sfGFP, and Tom20-HaloTag targeted to mitochondrial outer membrane fused by a P2A site (**Figure S5A**). HaloTag was labelled by TMR and transfected cells identified by sfGFP, and TMR signals were normalized to sfGFP intensities. We determined minor, but statistically significant reduced signal intensities for TMR-labelled Tom20-HT-SP5 compared to HaloTag and HaloTag M175Y (**Figure S5BC**).

To investigate how the three amino acid substitutions affect translocation by the T3SS, we analysed the thermal stability of the HaloTag and HT-SP5 protein. Therefore, protein denaturation was monitored via an increase in fluorescence of ROX, a dye that binds to hydrophobic residues that get exposed upon protein unfolding. The melting temperature of HaloTag and HT-SP5 was determined as 57 °C and 39 °C, respectively (**Figure 4CD**). Analyses using SYPRO Orange instead of ROX revealed comparable results (data not shown).

**Figure 4:**
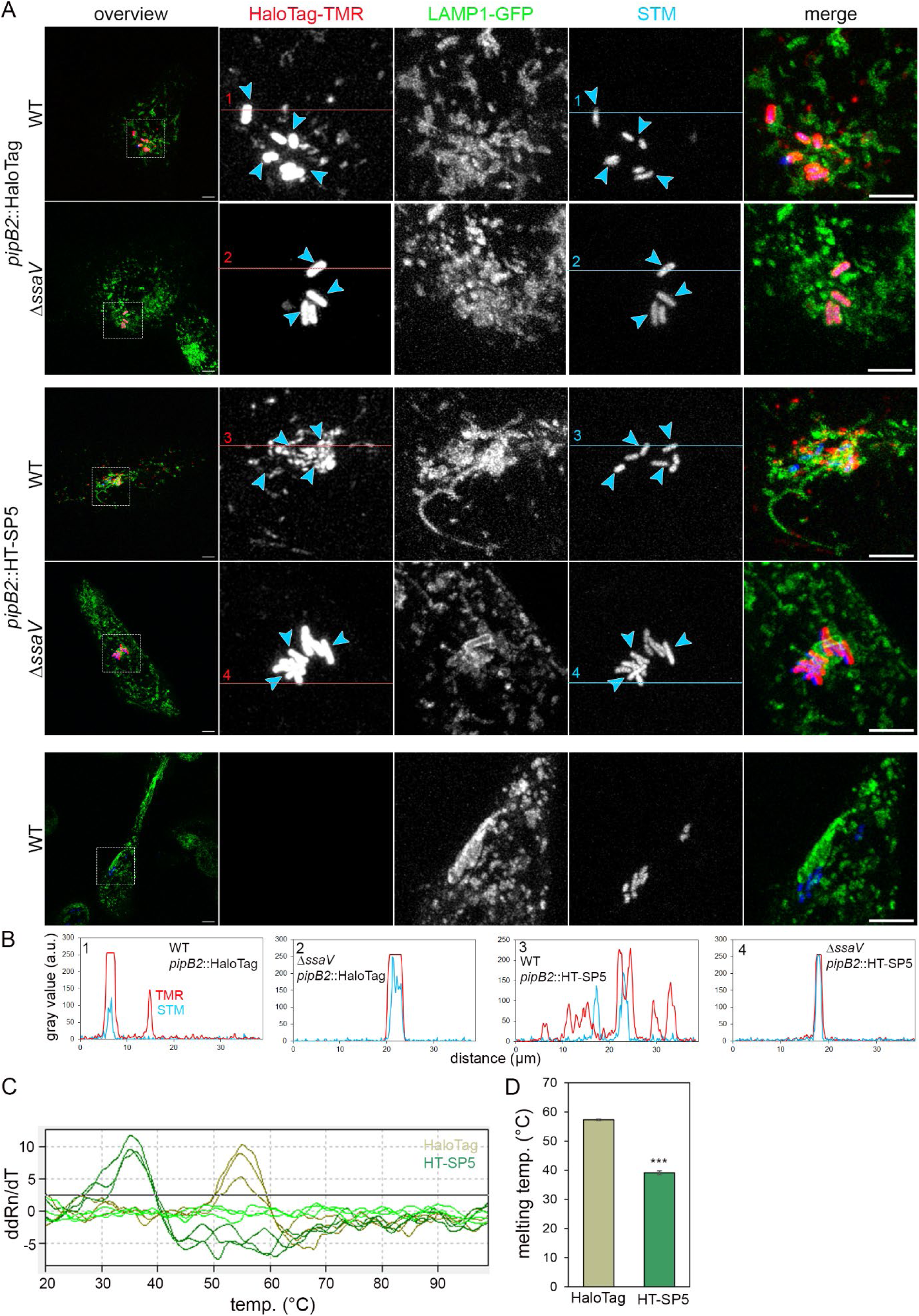
Distinct point mutations in HT-SP5 improve translocation of effector-HaloTag fusion proteins and reduce thermal stability compared to HaloTag. **A**) HeLa cells stably expressing LAMP1-GFP (green) were seeded in 8-well chamber slides and infection was performed at MOI 5 with STM WT, or MOI 20 with STM Δ*ssaV* expressing sfmTurquoise2ox (cyan) and PipB2-HaloTag fusions. Labelling reactions were performed directly before LCI was performed at 8 h p.i. using HTL-TMR (red) with a final concentration of 1 µM for 30 min at 37 °C. Scale bars, 10 µm. **B**) Line scans show signal intensities of translocated and intra-bacterial effector proteins in relation to the STM signal. **C**) Differential scanning fluorimetry of purified HaloTag and HT-SP5 proteins was performed using the dye ROX. Fluorescence intensities of the proteins were recorded during incubation ranging from 20 °C to 99 °C, and displayed as first derivative curves. **D**) Maximal temperatures corresponding to melting temperatures. Depicted are means and standard deviations of biological duplicates, with three technical triplicates each. Statistical significance was determined by unpaired, two-sided *t*-test as: ***, p < 0.001.

### Analysis of effector proteins at native expression levels

Translocation dynamics, subcellular localization and protein-interactions can be critically affected by the copy number of effector proteins. Hence, we investigated whether the improved translocation of our tagged effector protein would be sufficient to generate a detectable signal inside host cells without the need for plasmid-based overexpression required previously for many effector labelling approaches (Fritsch & Hensel, 2025). Instead of using HaloTag-effector fusions under control of the endogenous promoter encoded on low-copy plasmids, we used λ Red recombinase-mediated recombineering to create single-copy chromosomal HaloTag-fusions as reported previously (Barlag et al., 2016; Göser et al., 2019). Interestingly, slight overexpression of *sseJ*::HaloTag fusion using low copy number vectors resulted in the strong accumulation of SseJ effector proteins inside bacteria. This inhibition of effector translocation was less pronounced when effector-HaloTag-fusions were expressed in their chromosomal context. In contrast, the sub-cellular localization, distribution and effector-labelling intensity were indistinguishable between plasmid- and chromosomal-encoded *sseJ*::HT-SP5 fusions (**Figure S6**), indicating the possibility to analyse effector protein dynamics at native and manipulated effector levels.

### Applicability for T3SS effector proteins of other bacterial pathogens

To test the performance of HT-SP5 in analyses of T3SS effector proteins of other pathogens, we investigated enteropathogenic *E. coli* (EPEC). Of the approximately 20 T3SS effector proteins, the translocation dynamics and localization patterns of several effectors, including Map, are well known (Mills et al., 2013), and thus served us in our proof-of-concept analysis. The 203 amino acid effector protein Map harbours a mitochondrial targeting sequence at the N-terminus and a GTPase domain between residues 74-78, localizing Map to the mitochondria and plasma membrane, respectively, leading to disruption of mitochondrial functions and transient filopodia formation (Mandal et al., 2025). LCI of Lifeact-GFP expressing HeLa cells infected for 2 h by EPEC revealed clusters of adhering EPEC at F-actin-rich foci (**Figure S7**). We observed that HaloTag-labelled effector proteins were only poorly translocated, while the HT-SP5 fusion resulted in strong effector signals within host cells. These results indicate the general applicability of improved HaloTag for less interference with the T3SS translocation process to study effector dynamics in other pathogens.

### Improved performance of HT-SP5 in single molecule analyses of translocated effector proteins

Single-molecule localization and tracking microscopy (TALM) enables analyses of effector protein dynamics in host cells. We analysed whether the HaloTag variant HT-SP5 is suitable to follow PipB2 protein dynamics at the single molecule level (**Figure 6**). As previously shown, PipB2 fused to HaloTag can be localized and tracked on SIF 8 h p.i. or later (Göser et al., 2023). In contrast, PipB2 fused to HT-SP5 resulted in prominent trajectories on SIF tubules already 6 h p.i., when PipB2-HaloTag fusion trajectories were almost exclusively observed within STM. Enhanced intra-bacterial accumulation of PipB2-HT-SP5 was also not overserved at earlier time points, indicating rapid translocation upon expression, resulting in earlier onset of SIF formation as seen in the HeLa LAMP1-GFP overviews. The mobility of effector proteins was determined using pooled trajectories from mean square displacements (MSD) as a two-dimensional diffusion coefficient (DC) as reported previously (Göser et al., 2023). The movement of PipB2 fused to HaloTag and HT-SP5 on SIF membranes was similar (DC of 0.109 ± 0.03 and 0.102 ± 0.03), highlighting the applicability of HT-SP5 for TALM and effector protein dynamic studies.

### HT-SP5 enables analyses of effector protein translocation dynamics

Based on the improved translocation efficiency, we used HT-SP5 to follow the distribution of PipB2 as representative SPI2-T3SS effector protein over the course of STM infection (**Figure 5**). In the early phase (2 h p.i.), the signal intensity for PipB2-HT-SP5 was relatively low and most of the signals were associated with LAMP1-positive vesicles. At 3 h p.i., a prominent PipB2-HT-SP5 signal, mostly associated with LAMP-1 positive structures (vesicles, SCV and forming SIF membranes) was detectable. By LCI using cLSM, we followed the integration of vesicles double-positive for LAMP1 and PipB2-HT-SP5 into an extending network of SIF (**Figure 5C**). Compared to PipB2-HT-SP5, fusion with HaloTag delayed the observed effector dynamics by approximately 1 h (Göser et al., 2023), highlighting the feasibility of HT-SP5 for analyses of dynamics and kinetics of translocation and subcellular localization.

**Figure 5:**
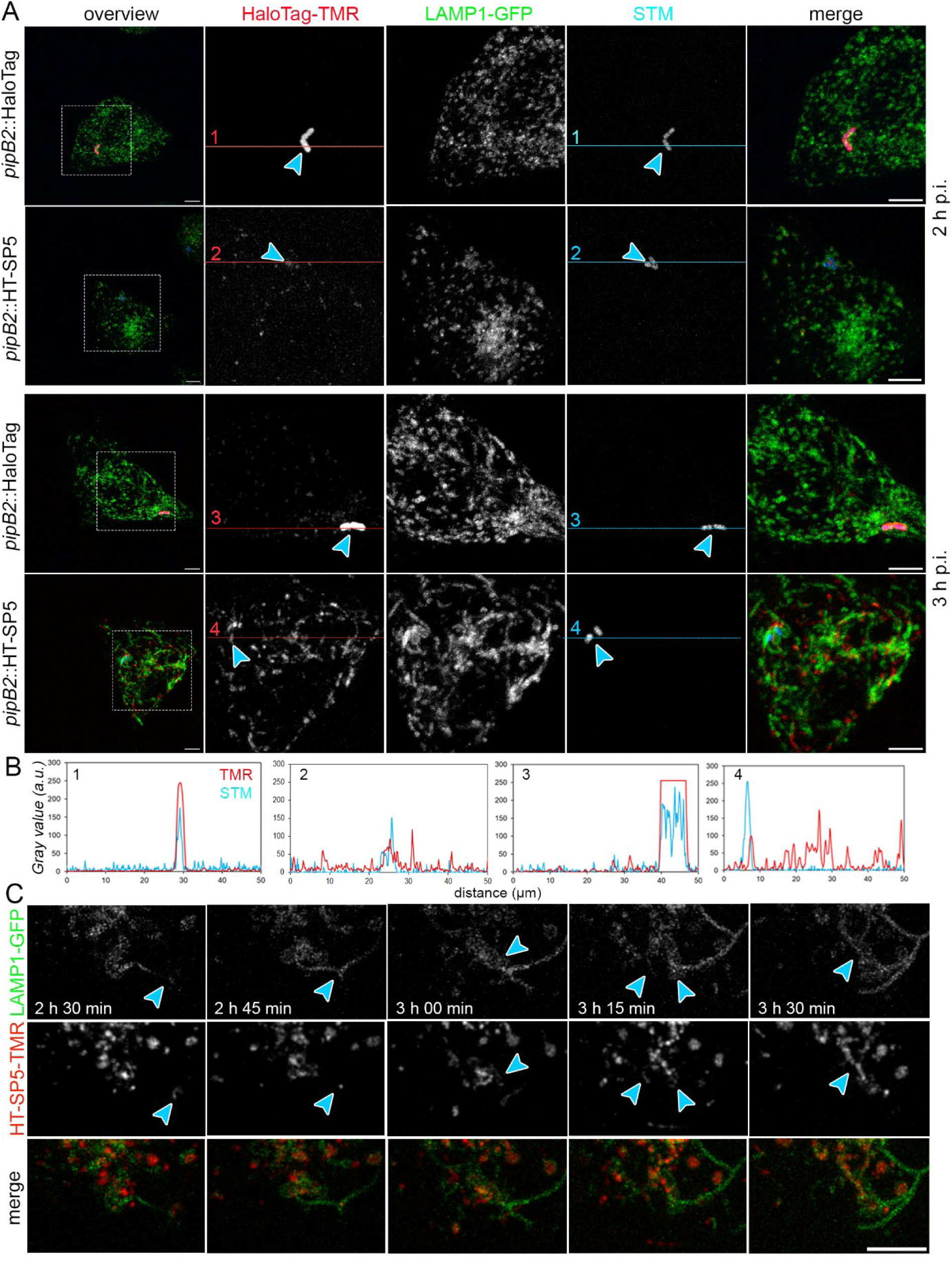
Effector fusions to HT-SP5 enable analyses of translocation kinetics early during infection. HeLa cells stably expressing LAMP1-GFP (green) were infected with STM WT expressing sfmTurquoise2ox (cyan) translocating PipB2 fused to HaloTag or HT-SP5. **A**) At 2 or 3 h p.i., cells were labelled with a final concentration of 1 µM HTL-TMR (red) before fixation and imaging. **B**) Line scans show signal intensities of translocated and intra-bacterial effector proteins in relation to the STM signal. **C**) Labelling reactions with HTL-TMR were performed directly before LCI at 2.5 h p.i. Scale bars, 10 µm.

Previously, pre-labelling of SopE-SNAP-tag enabled the visualisation of translocated effector proteins during bacterial invasion. However, the resulting signals were diffuse, making it impossible to identify single translocation events (Göser et al., 2019). To test the potential of effector fusion of HT-SP5 in LCI with high spatiotemporal resolution, we used Lattice Light Sheet microscopy (LLSM) and the fluorogenic HTL ligand MaP555 (Wang et al., 2020) to monitor translocated SPI1 effector proteins upon invasion of HeLa cells by STM (**Movie 1**, **Figure 7A**). At contact sites between STM and host cells, and SPI1-T3SS-induced actin cytoskeleton rearrangements, we detected distinct MaP555-labelled SopE-HT-SP5 signals at the poles of STM, followed by fast diffusion in host cells, and subsequent induction of membrane ruffling (**Figure 7Aiii**, 20 min p.i.). In host cells invaded at earlier time points, MaP555 signals were found located intracellular, clearly distant to the intracellular bacteria (**Figure 7Ai**, **Figure 7Aii**, **Movie 1**). In contrast, no signal was observed when using uninfected cells or STM WT as negative controls (**Movie 3AB**).

**Figure 6:**
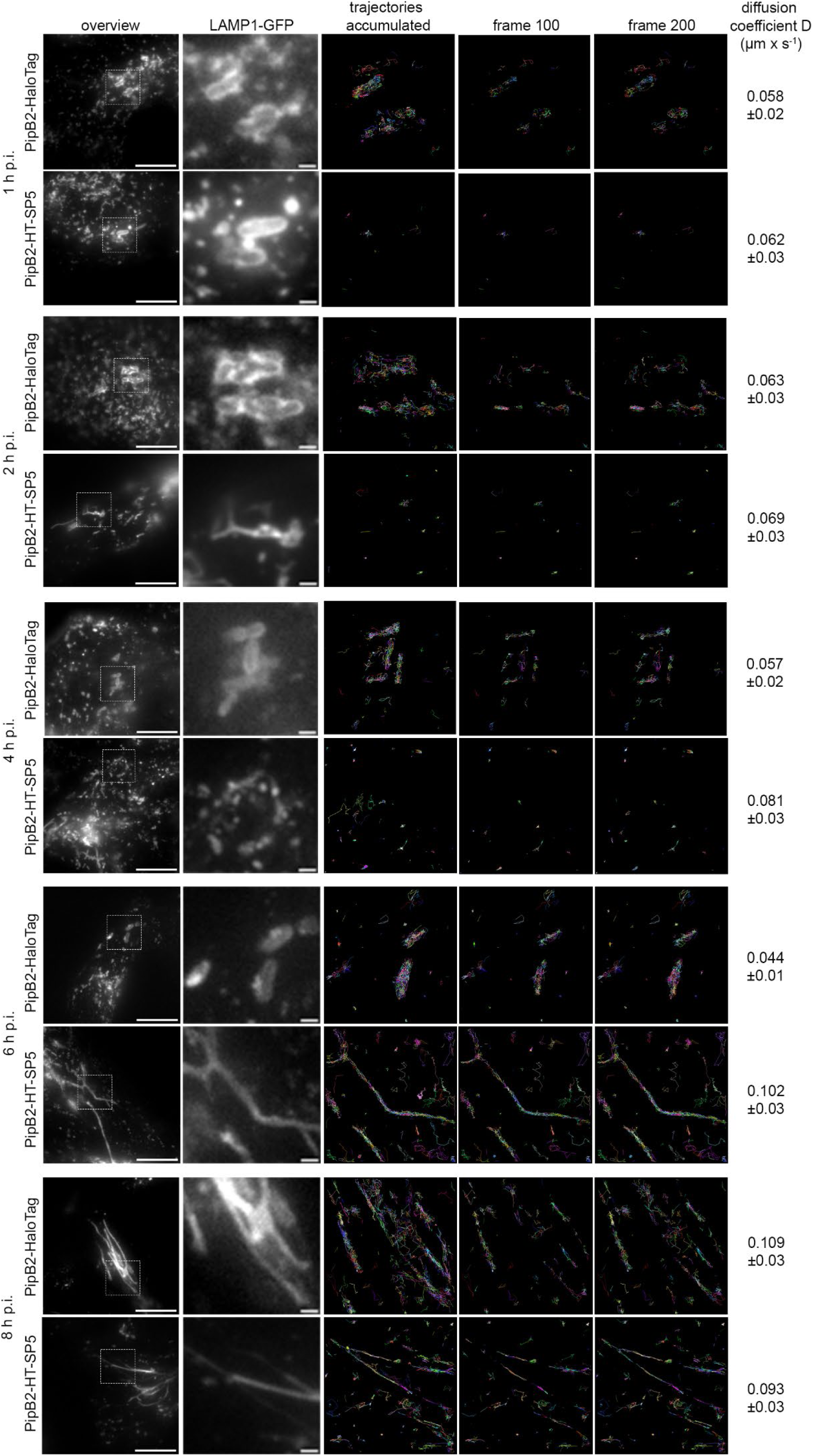
PipB2 fusion to HT-SP5 enables improved single molecule localization and tracking. For single molecule tracking, HeLa cells stably expressing LAMP1-GFP were infected at MOI 10 with STM Δ*pipB2* harbouring plasmids for expression of *pipB2*::HaloTag or *pipB2*::HT-SP5. After labelling with 20 nM HTL-TMR for 15 min at 37 °C, LCI was performed. Shown are representative images of LAMP1-GFP and trajectories for TMR signals (250 consecutive frames with a frame rate of 32 frames per s). The diffusion coefficient D of PipB2-HaloTag fusions was calculated using the Jaqaman algorithm based on at least 5 infected cells in two biological replicates. Scale bars, 10 and 1 µm in overviews and details, respectively.

**Figure 7:**
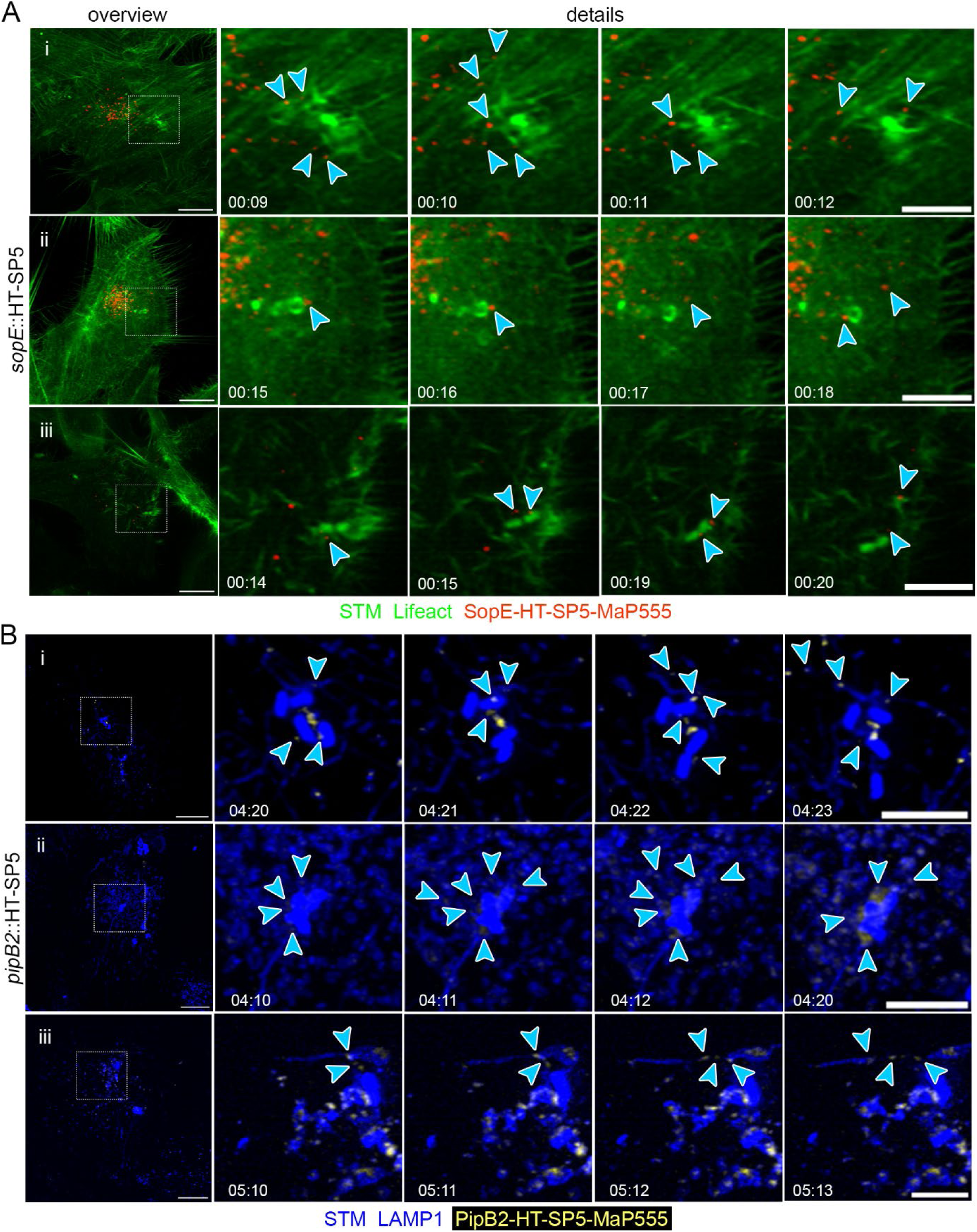
Fusions of effectors SopE or PipB2 to HT-SP5 enable analyses of dynamics of translocation and subcellular localization. **A**) HeLa cells stably expressing LifeAct-GFP (green) were infected by STM WT expressing mNeonGreen (green) and *sopE*::HT-SP5. Translocated effector was labelled by 20 nM fluorogenic MaP555 (red), and the diffusion of SopE-HT-SP5 was followed by LCI on a LLSM system. Three representative infected cells with translocation events are shown, see Movie 1 for kinetics of translocation and subcellular distribution. See **Movie 1** for time-lapse sequences, 3D projection, and details at higher magnification. **B**) HeLa cells stably expressing LAMP1-GFP (blue) were infected by STM WT harbouring plasmids for expression of mNeonGreen (blue) and *pipB2*::HT-SP5 under control of *tetR* P*tetA*. LCI was performed using LLSM in the presence of 20 nM fluorogenic MaP555 and 50 nM AHT induction of P*_tetA_*. Arrowheads indicate translocated PipB2-HT-SP5 labelled by MaP555 (yellow), and fusion of vesicles double-positive (white) for LAMP1-GFP and PipB2-HT-SP5 with the growing SIF network was followed in three representative infected cells. See **Movie 2** for kinetics of translocation and subcellular distribution, 3D projection, and details at higher magnification. Time points after infection are indicated by time stamp hh:mm. Scale bars, 10 µm (overview), 5 µm (detail).

Key challenges of the analysis of translocation of SPI2-T3SS effectors are the asynchronous expression and translocation, and heterogeneity of individual STM in host cells. To facilitate our analyses, we synchronised the expression of PipB2-HT-SP5 using an AHT-inducible promoter. This allowed us to identify at 4 h p.i. cells with an established SIF network as indicator for an active and functional SPI2-T3SS. Upon AHT induction, we observed the appearance of MaP555-labelled PipB2-HT-SP5 in close proximity to the intracellular bacteria (**Movie 2**, **Figure 7B**). Subsequent distribution in the cell either occurred along the SCV and SIF network or as vesicles double-positive for LAMP1 and PipB2-HT-SP5. As reported previously (Göser et al., 2023), we also detected fusion events of these vesicles with the growing SIF network. We did not observe PipB2-HT-SP5 freely diffusing into the host cytosol. In accordance with the lack of intra-bacterial signals, no signals were observed when using Δ*ssaV* mutants (**Movie 3B**), indicating that the used MaP555 concentration is not sufficient to cross the bacterial cell envelope and accumulate inside STM. Also, without AHT addition for *pipB2*-HT-SP5 expression, we did not detect any signal (**Movie 3C**), demonstrating the specificity of our approach.

These observations demonstrate the superior performance of HT-SP5 as tag for LCI analyses of T3SS translocation and the prospects to analyse early events of effector delivery into living host cells with high spatiotemporal resolution.

## Discussion

The elucidation of the molecular mechanisms and dynamics of translocation through T3SS is currently hampered by limited availability of T3SS-compatible reporter systems (Fritsch & Hensel, 2025). While NLuc and split NLuc-based assays allow kinetic analysis at high sensitivity and with a high dynamic range, they are restricted to population-based measurements, overlooking the bacterial heterogeneity inherent to host-pathogen interactions (Miao et al., 2024; Westerhausen et al., 2020). Conversely, the split-GFP assay enables the visualisation of translocated effector proteins within cells (Young et al., 2017), but its low sensitivity and slow complementation kinetics limit its utility for detecting early translocation events or performing detailed kinetic analyses. 4Cys-FlAsH has been used to study real-time kinetics, yet its application is constrained by non-specific labelling, cytotoxicity, and low sensitivity (Enninga et al., 2005; Van Engelenburg & Palmer, 2008). While genetic code expansion allows super-resolution imaging of effector proteins, low signal intensity and the requirement for two-photon fluorescence microscopy restrict this method to a number of specialised users (Singh & Kenney, 2024; Singh et al., 2021).

Here we describe an optimized HaloTag variant, HT-SP5, for biorthogonal labelling and tracking of effector proteins of bacterial T3SS. We demonstrated that effector-HT-SP5 fusions are efficiently translocated by invading and intracellular STM. In contrast to the previously used HaloTag, HT-SP5 does not interfere with translocation by T3SS, thereby preventing artifacts by intra-bacterial accumulation of effectors. Consequently, HT-SP5 allows detection and single-molecule tracking of translocated effector proteins several hours earlier than by using original HaloTag as reporter. This increased temporal sensitivity further allows the studying of the translocation dynamics of SPI1 and SP2-T3SS effector proteins. To our knowledge, this is the first study to identify SLE that are better suited for translocation by the T3SS, improving the toolkit for studying effector protein dynamics, especially effector translocation and the immediate post-translocation processes. While STM and EPEC have been used for validation in this work, applying HT-SP5 to other T3SS-deploying pathogens, or to other bacterial secretion systems, such as T4SS, should be the next step to confirm its general applicability.

Although our analyses revealed a 1.6-fold reduction in HTL-TMR labelling efficiency for HT-SP5 compared to HaloTag, this slight decrease in fluorescence intensity was fully compensated by the increased translocation efficiency. This was revealed by stronger effector signals within host cell cytoplasm and detection of effector proteins expressed by chromosomal genes. Previous research has shown that HaloTag labelling kinetics are highly substrate specific, varying by up to 6 orders of magnitude (Wilhelm et al., 2021). Interestingly, Miro-Vinyals *et al*., observed no differences in the fluorescence of both HaloTag variants when using the dye JF549 (Miro-Vinyals et al., 2021), indicating that alternative HaloTag ligands could further advance the analyses of effector protein dynamics. Application of the dimethylamino-styrylpyridium based dye F1, originally used to direct mutagenesis of HT-SP5 (Miro-Vinyals et al., 2021), resulted in high background staining, excluding this dye from our analysis (data not shown). Given the continuous development of new, improved substrates, future evaluation of the most suitable SLE dye is warranted. For example, recently Coïs *et al*. designed the fluorogenic ligand Red-Halo3, which exhibits 40% increased fluorescence *in vivo* and a reaction rate one order of magnitude higher with HaloTag M175Y V245A than with HaloTag *in vitro* (Coïs et al., 2024).

While we have excluded increased signal intensity as the cause for the elevated signal intensities for translocated effector proteins, we cannot rule out enhanced self-labelling efficiency with HTL-TMR as a contributing factor. Biochemical assays are required to determine labelling kinetics across different HaloTag substrates, especially TMR and MaP555. Our thermal stability assay indicates an enhanced compatibility of HT-SP5 with translocation by T3SS. Although recent studies highlighted that mechanical unfolding properties rather than thermal stability itself, are the determining factor (Akeda & Galan, 2005; DaPron et al., 2026; LeBlanc et al., 2021; Radics et al., 2014), a temperature difference of almost 20 °C likely affects the unfolding properties of the two HaloTag variants. We determined the melting temperature of HT-SP5 to be 39 °C, slightly above cell culture assay temperatures of 37 °C, indicating that HT-SP5 is more prone to unfolding required for T3SS translocation than the original HaloTag. To visualise SPI2-T3SS effector translocation despite the heterogeneity in T3SS activity, we synchronised PipB2 expression by a synthetically inducible system. While this enabled us to analyse the translocation process itself and track the timing between effector expression and translocation, it does not allow to understand the precise spatiotemporal regulation of SPI2-T3SS translocation activity and the hierarchy of effector translocation. This may be achieved by combining our approach with light-controlled activation of T3SS activity (Lindner et al., 2020), which would allow precise measurement of the interval between T3SS activation and subsequent effector translocation. To analyse effector hierarchy in more detail, our further studies will focus on multiplexed analyses combining SNAP-Tag and HT-SP5. Further approaches such as split-HaloTag (Wilhelm et al., 2025), or the use nanobody tags on effector proteins and host cell expression of cognate intrabodies (Fagbadebo & Rothbauer, 2022) may further complement SLE approaches.

## Conclusion

The investigation of the dynamic processes of effector proteins, including the timing and kinetics of effector protein translocation and their subcellular localization, that shape host-pathogen interactions, is still hampered by the limited toolbox to visualize effector proteins in living cells (Fritsch & Hensel, 2025). Here, we have introduced the mutated SLE tag HT-SP5 in combination with rhodamine-based ligands for analyses of effector proteins in single living cells. By introducing distinct point mutations in the HaloTag, we obtained a SLE that shows reduced thermal stability, resulting in an improved translocation efficiency upon fusion to T3SS effector proteins. This enables functional, temporal and spatial investigations of SPI1 and SPI2 effector proteins at native, physiological concentrations during host pathogen interactions. We demonstrated the possibility of cellular post-labelling and the application of fluorogenic dyes in no-wash experiments. We therefore expect that HT-SP5 will facilitate the analyses of the translocation process, such as activation and end of translocation, effector hierarchy, translocation kinetics and efficiency, effector dissemination, and cross-talk with other secretion systems.

In summary, this method provides a valuable tool for studying effector protein dynamics in individual living cells with single-molecule resolution and in multiplexing experiments combining various SLE tags.

## Acknowledgements

This work was supported by the Deutsche Forschungsgemeinschaft through projects P8 ‘Dynamic manipulation of functional plasticity of the host cell endosomal system by an intracellular pathogen’ (Project number 46752218) and Z2 ‘High-resolution imaging across spatiotemporal scales’ (Project number 467522186) in SFB1557. We thank Kay Johnsson, Max Planck Institute for Medical Research, Heidelberg, Germany, for the generous gift of MaP555; and gratefully acknowledge the expertise and advice from Rainer Kurre (iBiOs), and the technical support of Simon Schlichting, Ursula Krehe and Monika Nietschke.

## Materials and Methods

### Bacterial strains and growth conditions

Infection experiments were performed with *Salmonella enterica* serovar Typhimurium NCTC 12023 or SL1344 as WT strains, and isogenic mutant strains as specified in **Table 1**. For imaging, bacterial strains harboured p6739 or p6787 (**Table S 1**) for constitutive expression of sfmTurquoise2ox or mNeonGreen, respectively. Bacterial strains were routinely cultured in Luria broth (LB) containing 50 µg/ml carbenicillin or 12.5 µg/ml chloramphenicol if required to maintain plasmids. Construction of strains and recombinant DNA (**Table S1**) is described in detail in Suppl. Materials and Methods.

**Table 1.** Bacterial strains used in this study.

| Designation | relevant characteristics | reference |
| --- | --- | --- |
| *Salmonella enterica* serovar Typhimurium |  |  |
| NCTC12023 | Wild type strain | NCTC Colindale, lab stock |
| MvP1890 | $\Delta ssaV::FRT$ | (Noster et al., 2019) |
| MvP1944 | $\Delta pipB2::FRT$ | (Göser et al., 2023) |
| MvP1895 ($\Delta 5$) | $\Delta sopB::FRT \Delta sopA::FRT \Delta sopE2::FRT$<br>$\Delta sipA::FRT \Delta sopD::FRT$ | This study |
| MvP818 | $\Delta invC::FRT$ | (Gerlach et al., 2008) |
| MvP1764 | $\Delta invG::FRT$ | This study |
| MvP503 | $\Delta sifA::FRT$ | (Namakchian et al., 2018) |
| MvP2346 | *sseJ::HaloTag aph* | (Göser et al., 2019) |
| MvP3353 | *sseJ::HT-SP5 aph* | This study |
| Enteropathogenic *Escherichia coli* (EPEC) |  |  |
| E2348/69 | Wild-type strain, clinical isolate | Mathias Hornef (Aachen) |
| *Escherichia coli* |  |  |
| NEBα | general cloning strain | New England Biolabs |
| BL21 pLysS | expression of recombinant proteins | Lab stock |

### Labelling of HaloTag with TMR and MaP555 ligand

Mammalian cell lines used for infection are listed in **Table 2**, and maintenance and manipulation of cell lines is described in detail in Suppl. Materials and Method.

**Table 2:** Cell lines used in this study.

| Designation | relevant characteristics | reference |
| --- | --- | --- |
| HeLa | non-polarized epithelial cell | ATCC no. CCL-2 |
| HeLa LAMP1-GFP | stable expression of LAMP1-eGFP | (Krieger et al., 2014) |
| HeLa Lifeact-eGFP | for visualization of F-actin | (Scharte et al., 2024) |
| RAW264.7 | murine macrophage-like cell line | ATCC no. TIB-71 |

The self-labelling reaction of the mitochondrial marker Tom20 fused to HaloTag was performed in living cells at 37 °C 20 h after transfection of the cells with 150 nM HTL-TMR (ex. 545 nm, em. 575 nm) (Promega) in DMEM without 10% FCS for 30 min. Labelling of effector proteins fused to HaloTag was performed under the same conditions but with 1 µM HTL-TMR or 20 nM MaP555 unless stated otherwise. After labelling with HTL-TMR, the cells were washed 3-10 times with warm PBS and prepared for LCI, or were fixed in PFA for 20 min.

### (Live cell) fluorescence microscopy

For LCI, DMEM was replaced by imaging-medium consisting of Minimal Essential Medium (MEM) with Earle’s salts, 5 mM L-glutamine but without NaHCO_3_ and phenol red (Biochrom) supplemented with 30 mM HEPES (4-(2-hydroxyethyl)-1-piperazineethanesulfonic acid) (Sigma-Aldrich), pH 7.4. Imaging was performed by confocal laser-scanning microscopy (cLSM) using an SP5 system (Leica) equipped with an incubation chamber maintaining 37 °C and humidity during LCI. Image acquisition was done using a 40× (HCX PL APO CS 40×, NA 1.25–0.75) and 100× objectives (HCX PL APO CS 100×, NA 1.4–0.7) and the polychroic mirror TD 488/543/633 for the three channels GFP/ TMR/ DIC (Leica, Wetzlar, Germany). Additional, cLSM was performed using an Olympus FV-3000 system. The software LAS AF (Leica, Wetzlar, Germany) was used for setting adjustment and image acquisition.

### Lattice light sheet microscopy

Lattice Light-Sheet Microscopy (LLSM) was performed using a custom-build system based on the original design by the Eric Betzig group (Chen et al., 2014). Glass cover slips seeded with cells were inserted into a custom-built sample holder, ensuring that the sample was inserted at the correct position inside a sample bath, which contained Imaging medium with 10% FCS and 20 nM HTL-MaP555 at 37 °C. Stacks were acquired in sample-scan mode moving cells through a fixed light-sheet with a step size of 400 nm, which is equivalent to ∼217 nm slicing with respect to the Z-axis, when considering the sample scan angle of 32.8°. A dithered square lattice pattern was used, generated by multiple Bessel beams using an inner and outer numerical aperture of the excitation objective of 0.48 and 0.55, respectively. The fluorescent proteins mNeonGreen and sfGFP were excited via a 488 nm laser (2RU-VFL-P-300-488-B1R; MPB Communications Inc.), while the HTL-MaP555 dye was excited using a 561 nm laser (2RU-VFL-P-2000-561-B1R; MPB Communications Inc.), respectively. The final lattice light-sheet was generated by a water-dipping excitation objective (54–10–7 @488–910 nm, NA 0.66, Special Optics, NJ, USA), while emitted photons were collected by a water dipping detection objective (CFI Apo LWD 25XW, NA 1.1, Nikon, Tokyo, Japan). Emission was filtered using a 523/610 HC dual-band bandpass filter (Semrock) and imaged on an sCMOS camera (ORCA-Fusion, Hamamatsu, Japan) with a final pixel size of 103.5 nm and 23.5 ms exposure time. Raw data were further processed by usage of an open-source LLSM post-processing utility called LLSpy (https://github.com/tlambert03/LLSpy), which was used for de-skewing, deconvolution, channel registration, and transformation. Deconvolution was performed using experimental point spread functions, recorded from specific 100 nm sized FluoSpheres™ (Art. No. F8801 and No. F8803, ThermoFisher Scientific) and is based on the Richardson–Lucy algorithm using 10 iterations. Before acquisition, point spread functions were optimized using F8803 FluoSpheres™ and a custom-built adaptive optics system based on the AO KIT BIO from Imagine Optic. For channel alignment, registration and transformation, 200 nm sized fluorescent TetraSpeck™ microspheres (Art. No. T7280, ThermoFisher Scientific) were imaged with a step size of 350 nm. Chanel transformation was done with a 2-step transformation method that deploys an affine transformation in xy and a rigid transformation in z.

### Quantification of fluorescence

The ratio of intracellular HTL-conjugated dyes bound to Tom20-HaloTag to cytosolic sfGFP was analysed via the software FIJI by means of sum projections of Z stacks of whole cells acquired over the same z-range. The mean intensity of each labelled area (TMR, sfGFP) was subtracted from the mean intensity of an unlabelled area (background). The cell was segmented using a threshold calculated by applying the Huang method and the mean intensities for both the sfGFP an TMR channels were obtained. To quantitatively compare effector fusions to HaloTag and HT-SP5, line scan analysis of maximal intensity projections showing TMR, LAMP1-GFP, or sfmTurquoise2ox signals were performed using Image J/Fiji (1.54p).

### Quantification of SCV and SIF network volume

For the analysis of the SCV and SIF volumes in relation to the volume of the bacteria, HeLa cells were infected at MOI 10 with *Salmonella* WT or Δ*pipB2* strain expressing sfmTurquoise2ox and *pipB2* fused to HaloTag::HA or HT-SP5::HA as described. LCI was performed 8 h p.i. The volume of intracellular bacteria and the interconnected SCV and SIF network were determined for at least 30 cells per condition using the three-dimensional reconstruction tool of Imaris (Bitplane, Zürich, Switzerland). As reported previously, SIF and STM volumes were adjusted using auto-threshold and smoothing of 0.3 (Namakchian et al., 2018).

### Differential scanning fluorimetry

For expression and purification of His-tagged HaloTag and HT-SP5, *E. coli* BL21 pLysS strains harbouring p6300 or p6303 were subcultured in LB at 37 °C until OD_600_ of 0.6, followed by the addition of 50 ng/ml AHT and incubation for 3 h at 30 °C. Bacterial cells were harvested by centrifugation, and pellets were disrupted by sonication in 20 mM sodium phosphate, pH 7.4, 500 mM NaCl, 50 mM imidazole containing protease inhibitors (Roche), lysozyme, and DNAse. The lysate was centrifuged for 1 h at 50,000 x *g* at 4 °C and recombinant His-tagged proteins were purified using His Spin Trap columns (GE Healthcare) according to the manufacturers’ instructions. Fusion proteins were eluted in a buffer containing 20 mM sodium phosphate, pH 7.4, 500 mM NaCl and 500 mM imidazole. The final purification step encompassed a size exclusion chromatography run (Superdex 75 Prep-Grade, GE Healthcare) using a buffer with 20 mM sodium phosphate, pH 7.4, 500 mM NaCl.

Protein stability was determined as described previously using differential scanning fluorimetry (Niesen et al., 2007). Purified protein was used at a concentration of 1 µg/µl and mixed with buffer, SYPRO Orange (Fisher 11510746) and ROX (Thermo 4461146) according to recommendations of the manufacturer. Temperature was increased every minute by 1 °C until reaching 99 °C and fluorescence intensity was recorded using the qTOWER3 touch. For subsequent analyses, the qPCR soft 4.1 software (Analytik Jena AG) was used.

### Statistics

Each experiment was repeated at least twice with similar results, using independent experimental samples. Statistical analysis was performed using the Students’ unpaired two-tailed *t*-test.

## Notes

### Competing Interest Statement

The authors have declared no competing interest.

