## Supplementary material for "Improved HaloTag for analyses of translocation of type III secretion system effector proteins": Suppl. Materials

### Supplementary Materials and Methods

*Generation of plasmids*

Plasmids used in this study (**Table S1**) were constructed using primers listed in **Table S2** to generate DNA fragments specified in **Table S3**. Fragments were used for modular designed Gibson Assembly. Plasmid p6469 with chloramphenicol resistance cassette was used as basis vector for the generation of fluorescent plasmids p6739 and p6787. The *ompA* promoter and sfmTurq2ox were amplified from p6724 (pACYC $P_{ompA}$::sfGFP) and p5384 (pWRG167_sfmTurquoise2ox STM), respectively, while the mNeonGreen fragment was obtained as codon-optimized synthetic DNA fragment (IDT).

For generation of mutant alleles of HaloTag, site-directed mutagenesis (SDM) was performed using the Q5 mutagenesis kit (NEB) basically according to manufacturers' instructions. Plasmids p6293 and p6300 were used as templates for amplification with SDM primers (IDT) listed in **Table S2**. Multiple exchanges were generated by repeated rounds of SDM, and resulting plasmids were confirmed by DNA sequencing (Microsynth). Confirmed HaloTag-HA alleles were PCR-amplified using primer set 3f-link-HaloTag and 3r HA-pWSK. To generate low copy number plasmids (**Table S1**) with various combinations of effector genes and HaloTag alleles, Gibson assembly with modules V p5943, 1 (promoter modules), 2 (effector module), or 1+2 (promoter+effector module), and 3 (HaloTag module) as described in **Table S3** was performed.

For generation of chromosomal fusions with HaloTag::HA alleles, λ Red recombineering was used basically as described (Barlag et al., 2016; Gerlach et al., 2007). Plasmid p6861 was generated as template vector for generation of HT-SP5::HA cassettes as targeting DNA, with primer set sseJ-GS RedIn For and sseJ RedIn Rev for generation of a 3’ fusion to *sseJ*.

*Transient transfection of cells*

HeLa cells were cultured for one day and transfected using FuGENE HD reagent (Promega) according to manufacturer's instruction. Plasmids used for transfection are listed in **Table S1**. Briefly, 0.5 µg of plasmid DNA were dissolved in 25 µl OptiMEM without iFCS and mixed with 1 µl FuGENE reagent (ratio of 1:2 for DNA to FuGENE). After 10 min incubation at room temperature (RT) the transfection mix was added to the cells in DMEM with 10% iFCS for at least 19 h. Before staining, cells were provided with fresh medium without transfection mix for 1 h. Labelling was performed using 150 nM HTL-TMR for 30 min. Fixed cells were imaged as described above. Intensity measurements of GFP and TMR channel were performed after Z-stack slices were summed, the background was subtracted and the cell segmented using ImageJ 1.53t.

*Host cell infection*

For the infection of RAW264.7 cells, overnight cultures of *Salmonella* strains were used, while for infection of HeLa cells, overnight cultures were diluted 1:31 in fresh LB, and subcultured for 3.5 h. Infection was performed at indicated MOI for 25 min, after centrifugation for 5 min at 500 x g to synchronize infection. Subsequently, cells were washed thrice with prewarmed phosphate-buffered saline (PBS) and incubated for 1 h with medium containing 100 mg/ml gentamicin (Applichem) to kill extracellular bacteria. Medium was replaced by medium containing 10 mg/ml gentamicin for the rest of the experiment and cells were incubated et 37 °C in a humidified atmosphere of 5% $CO_2$.

Intracellular replication was determined using gentamicin protection assays. In brief, $2x10^5$ RAW264.7 cells, seeded at least 24 h prior to infection in 24-well plates, were infected in triplicates at indicated MOI. Cell lysis was performed 2 and 16 h p.i. after washing thrice with PBS by incubation with PBS containing 0.1% Triton X-100 for 15 min. Colony forming units (CFU) were determined by spotting serial dilutions of the lysates onto Mueller-Hinton (MH) agar plates. Phagocytosis was determined as percentage of inoculum internalized, and intracellular replication as calculated as ratio of CFU recovered at 16 h and 2 h p.i.

EPEC strains were cultured overnight in in LB broth containing 50 µg/ml carbenicillin at 37 °C with agitation in a roller drum. On the day of infection, a static subculture with ratio of 1:51 was grown at 37 °C in DMEM (4.5 g/l glucose, 3.7 g/l $NaHCO_3$, containing stable L-glutamine and Na-pyruvate) without iFCS. After 3 h, $OD_{600}$ was measured and the subculture was diluted to $OD_{600}$ 0.2 in 1 ml PBS. Cells were infected with MOI of 30 for 3 h before imaging.

*Invasion assay*

HeLa cells were seeded in surface-treated 24-well plates (TPP) at least 24 h prior to infection to reach confluency (ca. $2x10^5$ cells). Cells were infected as described above at MOI 5. Serial dilutions of inoculum and lysates at 1 h p.i. were spotted onto MH agar. The percentage of internalized bacteria was calculated.

*Detection of secreted effector proteins by Western blotting*

Overnight cultures of STM strains were subcultured (1:31) in LB broth at 37 °C and harvested after 2, 4, and 6 h. The filtered supernatant was precipitated overnight at 4 °C with 10% trichloroacetic acid. The pellets of precipitated supernatants were washed twice with acetone and air dried. Total bacterial cells and precipitated secreted proteins were adjusted based on $OD_{600}$ and dissolved in SDS-PAGE loading buffer containing 25% glycerol, 4% SDS, 50 mM Tris, 2% Mercaptoethanol, pH 6.8, and boiled for 2 min. 10 µl of samples were loaded onto a 10% SDS-PAGE gel. After electrophoresis, samples were blotted onto a 0.45 µm nitrocellulose membrane using a semidry electrophoretic transfer unit (Bio-Rad). Blots were incubated with a primary antibody directed against the HA epitope tag (1:10,000) and a secondary antibody anti-rat IgG antibody conjugated to horseradish peroxidase (HRP) (1:10,000). The detection was performed using the ECL detection kit (Pierce) and Chemidoc imaging system (Bio-Rad).

*Quantification of effector protein secretion by flow cytometry analyses*

For subsequent flow cytometry analyses using a Attune NxT (Thermo Fisher), bacteria were sub-cultured for 3.5 h in LB and then adjusted to $OD_{600}$ of 0.04 with PBS. Labelling was performed using 1 µM TMR for 40 min at 37 °C and 800 rpm. Unbound TMR was removed by washing the bacteria thrice with PBS. At least 25,000 bacteria-sized particles were analysed by flow cytometry in three independent experiments each, and TMR fluorescence was determined using a flow rate of 12.5 µl/min and the YL1 filter. Data were analysed with Attune NxT Software.

*Quantification of effector protein translocation by fluorometry*

Overnight cultures of STM expressing *pipB2*::HaloTag or *pipB2*::HT-SP5 were used to infect RAW264.7 macrophages at MOI 5. Effector-HaloTag fusions were labelled 8 h p.i. using 1 µM HTL-TMR for 30 min at 37 °C. Cells were washed 7 times to remove unbound ligand, and subsequently detached or lysed using Triton X-100. The cellular and bacterial suspension was pelleted, resuspended in PBS, and a total of 200 µl was then added to each well of the microplate. Fluorescence intensity was measured directly using an excitation and emission wavelength of 540 nm (±9 nm) and 570 nm (±20 nm), respectively, by a microplate fluorescence reader (Infinite 200 Pro M-Plex, Tecan). RAW264.7 macrophages infected with STM WT was used as reference.

### Supplementary Tables

**Table S 1: Plasmids used in this study**

| Plasmid | relevant genotype | reference |
|---|---|---|
| pWSK29 | low copy number vector, $Amp^R$ | lab stock |
| pWRG167 | $P_{EM7}$::sfGFP | Lab stock |
| pLX304 | lentiviral transfection vector | Addgene |
| p2104 | $P_{\mathit{sifA}}$ *sifA*::M45 | (Hansen-Wester et al., 2002) |
| p2795 | template plasmid for λ Red recombineering, $Amp^R$, $Kan^R$ | (Gerlach et al., 2007) |
| p3773 | pWSK29 *tetR* $P_{\mathit{tetA}}$ | (Hansmeier et al., 2017) |
| p4042 | $P_{\mathit{sopB}}$ *sopB*::HA | (Felipe-Lopez et al., 2023) |
| p4043 | $P_{\mathit{sopE}}$ *sopE*::HA | (Felipe-Lopez et al., 2023) |
| p4115 | $P_{\mathit{sipA}}$ *sipA*::L16::HaloTag::HA | (Göser et al., 2019) |
| p4117 | $P_{\mathit{sopE}}$ *sopE*::L16::HaloTag::HA | (Göser et al., 2019) |
| p4286 | $P_{\mathit{sseJ}}$ *sseJ*::L16::HaloTag::HA | (Göser et al., 2019) |
| p4295 | $P_{\mathit{pipB2}}$ *pipB2*::L16::HaloTag::HA | (Göser et al., 2019) |
| p4305 | $P_{\mathit{sifA}}$ *sifA*::L16::HaloTag::HA | (Göser et al., 2019) |
| p5384 | pWRG167_sfmTurquoise2ox STM | This study |

| | | |
|---|---|---|
| p5665 | pLX304 Tom20::HaloTag | (Liss et al., 2015) |
| p5943 | pWSK29 with T7 transcriptional terminator (TT) | (Hensel, 2026) |
| p6229 | $P_{pipB2}$ *pipB2*::HaloTag::HA M175Y V245A | This study |
| p6293 | $P_{pipB2}$ *pipB2*::HaloTag::HA | This study |
| p6300 | *tetR* $P_{tetA}$::H6:: HaloTag::HA | This study |
| p6303 | *tetR* $P_{tetA}$::H6::HaloTag::HA M175Y V249A L271D | This study |
| p6469 | pACYC184 ins p5943 MCS and TT | (Hensel, 2026) |
| p6600 | $P_{sopB}$ *sopB*::HaloTag::HA M175Y V249A L271D | This study |
| p6601 | $P_{sopE}$ *sopE*::HaloTag::HA M175Y V249A L271D | This study |
| p6604 | $P_{sseJ}$ *sseJ*::HaloTag::HA M175Y V249A L271D | This study |
| p6605 | $P_{sifA}$ *sifA*::HaloTag::HA M175Y V249A L271D | This study |
| p6615 | $P_{sifA}$ *sifA*::HaloTag::HA M175Y | This study |
| p6611 | $P_{sopE}$ *sopE*::HaloTag::HA M175Y | This study |
| p6620 | $P_{sopB}$ *sopB*::HaloTag::HA | This study |
| p6621 | $P_{sopE}$ *sopE*::HaloTag::HA | This study |
| p6625 | $P_{sifA}$ *sifA*::HaloTag::HA | This study |
| p6678 | $P_{pipB2}$ *pipB2*::HA | This study |
| p6679 | $P_{pipB2}$ *pipB2*::HaloTag::HA M175Y V249A L271D | This study |
| p6680 | $P_{pipB2}$ *pipB2*::HaloTag::HA M175Y | This study |

| | | |
|---|---|---|
| p6787 | $P_{ompA}$::mNeonGreen | This study |
| p6724 | pACYC $P_{ompA}$::sfGFP | This study |
| p6739 | $P_{ompA}$::*s*fmTurquoise2ox | This study |
| p6861 | p2795 with HT-SP5::HA, template plasmid | This study |
| p7101 | $P_{pipB2}$ *pipB2*::HaloTag::HA M175Y V249A L271D F144L | This study |
| p7102 | $P_{pipB2}$ *pipB2*::HaloTag::HA I211V | This study |
| p7138 | pLX Tom20::HaloTag::HA M175Y::P2A::sfGFP | This study |
| p7156 | pLX Tom20::HaloTag::HA::P2A::sfGFP | This study |
| p7157 | pLX Tom20::HaloTag::HA M175Y V245A L271D::P2A::sfGFP | This study |
| p7169 | $P_{pipB2}$ *pipB2*::HaloTag::HA M175Y V249A L271D I211V | This study |
| p7175 | $P_{map}$ *map*::HaloTag::HA | This study |
| p7178 | Pmap map::HaloTag::HA M175Y V249A L271D | This study |
| p7198 | PsicA sipA::HaloTag::HA M175Y V249A L271D | This study |
| p7200 | PsicA sipA::HaloTag::HA | This study |
| p7213 | tetR PtetA::pipB2::HaloTag::HA M175Y V249A L271D | This study |

**Table S 2: Oligonucleotides used in this study:**

| Designation | sequence (5’ – 3’) | internal # |
|---|---|---|
| Cloning | | |
| Vr-pWSK | ATGGCCAGAAgtgGCCCGCGAGA | 93/21 |
| Vf-pWSK-Pnah | CAGCTTTTGTTCCCTTTAGTGA | 85/09 |
| Vf-pWSK29 | GAATTCCTGCAGCCCGGGG | 61/04 |
| Vr p5536 | GGATCCCCATCGATCCTTATCGT | 122/01 |
| Vf-pLX304 | TTCTTGTACAAAGTGGTTGGTAA | 88/47 |
| Vr-pLX304 | TGATCCCGACAGTTAGCC | 88/48 |
| 1f pWSK-tetR | CCCACTACGTGAACCATCAAACAAAGTTCCTATACTTTCTAGAGAATAGG | 129/56 |
| 1r PtetA-Plink | CTGGATGGATGTAGCATGTTCACTTTTCTCTATCACTGATAGGG | 129/60 |
| 1f pWSK-PompA | CCCACTACGTGAACCATCAAACGACAGCATTCCGGGCTAAAAATTC | 146/35 |
| 1r PompA-link | CTGGATGGATGTAGCATGCCAAAATACGCCATGAATATCTCC | 146/34 |
| 1f pWSK-PsicA | CCCACTACGTGAACCATCAAACGTTATGATCTGGTGAGTCTGG | 152/17 |
| 1r PsicA-link | CTGGATGGATGTAGCATGTCACCGACTTTGTAGAACTTAAC | 152/16 |
| 1f pWSK-PsifA | CCCACTACGTGAACCATCAAACCGCTAACAAATCCACACGC | 136/14 |
| 2r sifA-link | GGATCCGGAACCGCTTCCTAAAAAACAACATAAACAGCCGC | 141/62 |
| 1f pWSK-PpipB2 | CCCACTACGTGAACCATCAAACTAATAAAATGCCTGAACACG | 141/65 |
| 2r pipB2-link | GGATCCGGAACCGCTTCCAATATTTTCACTATAAAATTCG | 141/60 |
| 1f pWSK-PsopB | CCCACTACGTGAACCATCAAACTGGGTTTTCAATAAAAGTTG | 143/78 |
| 2r2 sopB-link | GGATCCGGAACCGCTTCCAGATGTGATTAATGAAGAAATGC | 146/76 |
| 1f pWSK-sopE | CCCACTACGTGAACCATCAAACCTTTGGACGCCTGCCACC | 142/73 |
| 2r sopE-link | GGATCCGGAACCGCTTCCGGAGGCATTCTGAAGATACTTATTC | 142/78 |
| 1f pWSK-Pmap (EPEC) | CCCACTACGTGAACCATCAAACAGATCTTTGCAAAATTGTTCATTC | 158/70 |
| 2f2 map-link (EPEC) | GGATCCGGAACCGCTTCCCAGCCGAGTATCCTGCACATTGTC | 159/27 |
| 2f link-pipB2 | CATGCTACATCCATCCAGCTGTCTCTGGGAGAAAATATATG | 141/59 |
| 2f link-sipA | CATGCTACATCCATCCAGAACAGAAGAGGATATTAATAATGG | 144/01 |
| 2r sipA-link | GGATCCGGAACCGCTTCCACGCTGCATGTGCAAGCCATCAAC | 146/77 |
| 1f PtetA-Halo | ATAAGGATCGATGGGGATCCGGATCCGAAATCGGTACT | 125/67 |
| 1r Halo-pWSK | CCCGGGCTGCAGGAATTCCTAACCGGAAATCTCCAG | 125/68 |
| 2f link-RBS_mNeon | CATGCTACATCCATCCAGTGAGAAAGAGGAGAAAAGTATGGTGTCCAAGGGTGAGGAGG | 150/05 |

| | | |
|---|---|---|
| 3r mNeon_stop-pWSK | CTAAAGGGAACAAAAGCTGTTACTTGTAAAGCTCATCCATGCCC | 150/12 |
| 2f link-RBS_sfmTurq2ox | CATGCTACATCCATCCAGTGAGAAAGAGGAGAAAAGTATGGTTTCCAAAGGAGAGGAG | 149/42 |
| 3r sfmTurq2ox-pWSK | CTAAAGGGAACAAAAGCTGTTATTTGTACAACTCATCCATAC | 149/44 |
| 3f link-HaloTag | GGAAGCGGTTCCGGATCCGAAATCGGTACTGGCTTTCCATTC | 141/10 |
| 3r HA-pWSK | CTAAAGGGAACAAAAGCTGTTAAGCGTAGTCTGGGACGTC | 141/19 |
| 1f link-HaloTag | GGGATCCACCGGTCGCCACCGAAATCGGTACTGGCTTTCCATTC | 126/12 |
| 1r HA-P2A | TCTCCAGCCTGCTTCAGCAGGCTGAAGTTAGTAGCAGCGTAGTCTGGGACGTCGTATGGG | 158/42 |
| 2f pLX-Tom20 | CTGGCTAACTGTCGGGATCAGCCACCATGGTGGGTCGGAACAGCGCC | 126/23 |
| 2r Tom20-link | GTGGCGACCGGTGGATCCCGGATTTCCACATCATCTTCAGCC | 126/24 |
| 3f P2A-sfGFP | GAAGCAGGCTGGAGACGTGGAGGAGAACCCTGGACCTCGCAAAGGCGAAGAACTGTTTAC | 158/41 |
| 1r sfGFP-pLX304 | CCAACCACTTTGTACAAGAATTATTTATACAGTTCATCCATG | 126/14 |

<u>λ Red recombineering</u>

| | | |
|---|---|---|
| sseJ-GS RedIn For | AATGTTAGAAAGTTTTATAGCTCATCATTATTCCACTGAAGGAAGCGGTTCCGGATCC | 152/78 |
| sseJ RedIn Rev | GCTGTGTTTTGCTCAAGGCGTACCGCAGCCGATGGAACTTGTAGGCTGGAGCTGCTTCGA | 152/79 |

<u>SDM</u>

| | |
|---|---|
| HaloTag M175Y For | TACGCTGCCGtatGGTGTCGTCC |
| HaloTag M175Y Rev | CCCTCGATAAAAACGTTC |
| HaloTag V245A For | CACCCCAGGCgctCTGATCCCAC |
| HaloTag V245A Rev | CCCCAGAACAGCAGCTTCGG |
| HaloTag L271D For | CGGCCCGGGTgatAATCTGCTGC |
| HaloTag l271D Rev | ATGTCCACAGCCTTGCAG |
| HaloTag F144L For | ATGGCCAGAAgtgGCCCGCGAGA |
| HaloTag F144L Rev | TCGTCCCAGGTCGGGATAG |
| HaloTag I211V For | CGAGCTGCCAgtgGCCGGTGAGC |
| HaloTag I211V Rev | TTTGGGAAGCGCCACAGTG |

**Table S 3: Gibson assembly fragments used in this study**

| Fragments | template | for primer | rev primer |
|---|---|---|---|
| V p5493 | p5943 | Vr-pWSK | Vf-pWSK-Pnah |
| V p5536 | p5536 | Vf-pWSK29 | Vr p5536 |
| V pLX304 | pLX304 | Vf-pLX304 | Vr-pLX304 |
| 1 tetR PtetA-link | p3773 | 1f pWSK-tetR | 1r PtetA-Plink |
| 1 PompA-link | STM genomic | 1f pWSK-PompA | 1r PompA-link |
| 1 PsicA-link | STM genomic | 1f pWSK-PsicA | 1r PsicA-link |
| 1+2 PsifA sifA-link | STM genomic | 1f pWSK-PsifA | 2r sifA-link |
| 1+2 PsopE sopE-link | STM genomic (SL) | 1f pWSK-PsopE | 2r sopE-link |
| 1+2 PpipB2 pipB2-link | STM genomic | 1f pWSK-PpipB2 | 2r pipB2-link |
| 1+2 PsopB sopB-link | STM genomic | 1f pWSK-PsopB | 2r2 sopB-link |
| 1+2 PmapA mapA-link | EPEC genomic | 1f pWSK-Pmap (EPEC) | 2r2 map-link |
| 2 pipB2-link | STM genomic | 2f link-pipB2 | 2r pipB2-link |
| 2 sipA-link | STM genomic | 2f link-sipA | 2r2 sipA-link |
| 2 link-mNeonGreen | synthetic | 2f link-RBS_mNeon | 3r mNeon_stop-pWSK |
| 2 link-sfmTurquoise2ox | p5384 | 2f link-RBS_sfmTurq2ox | 3r sfmTurq2ox-pWSK |
| 3 link-HaloTag-HA | p4295 | 3f link-HaloTag | 3r HA-pWSK |
| 3 link-HaloTag-HA M175Y | p6680 | 3f link-HaloTag | 3r HA-pWSK |
| 3 link-HaloTag-HA I211V | p7102 | 3f link-HaloTag | 3r HA-pWSK |
| 3 link-HaloTag-HA M175Y V245A | p6229 | 3f link-HaloTag | 3r HA-pWSK |
| 3 link-HaloTag-HA M175Y V249A L271D | p6679 | 3f link-HaloTag | 3r HA-pWSK |
| 3 link-HaloTag-HA M175Y V249A L271D F144L | p7101 | 3f link-HaloTag | 3r HA-pWSK |
| 3 link-HaloTag-HA M175Y V249A L271D I211V | p7169 | 3f link-HaloTag | 3r HA-pWSK |
| 1 link-6H::HaloTag | p3780 | 1f PtetA-Halo | 1r Halo-pWSK |
| 1 link-6H::HT-SP5 | p6679 | 1f PtetA-Halo | 1r Halo-pWSK |
| 1 HaloTag-HA-P2A | p4295 | 1f link-HaloTag | 1r HA-P2A |
| 1 HaloTag-HA-P2A M175Y | p6680 | 1f link-HaloTag | 1r HA-P2A |
| 1 HaloTag-HA-P2A M175Y V249A L271D | p6679 | 1f link-HaloTag | 1r HA-P2A |
| 2 pLX-Tom20 | p5665 | 2f pLX-Tom20 | 2r Tom20-link |
| 3 P2A-sfGFP-pLX | pWRG167 | 3f P2A-sfGFP | 1r sfGFP-pLX304 |

### Suppl. Figures and Figure Legends

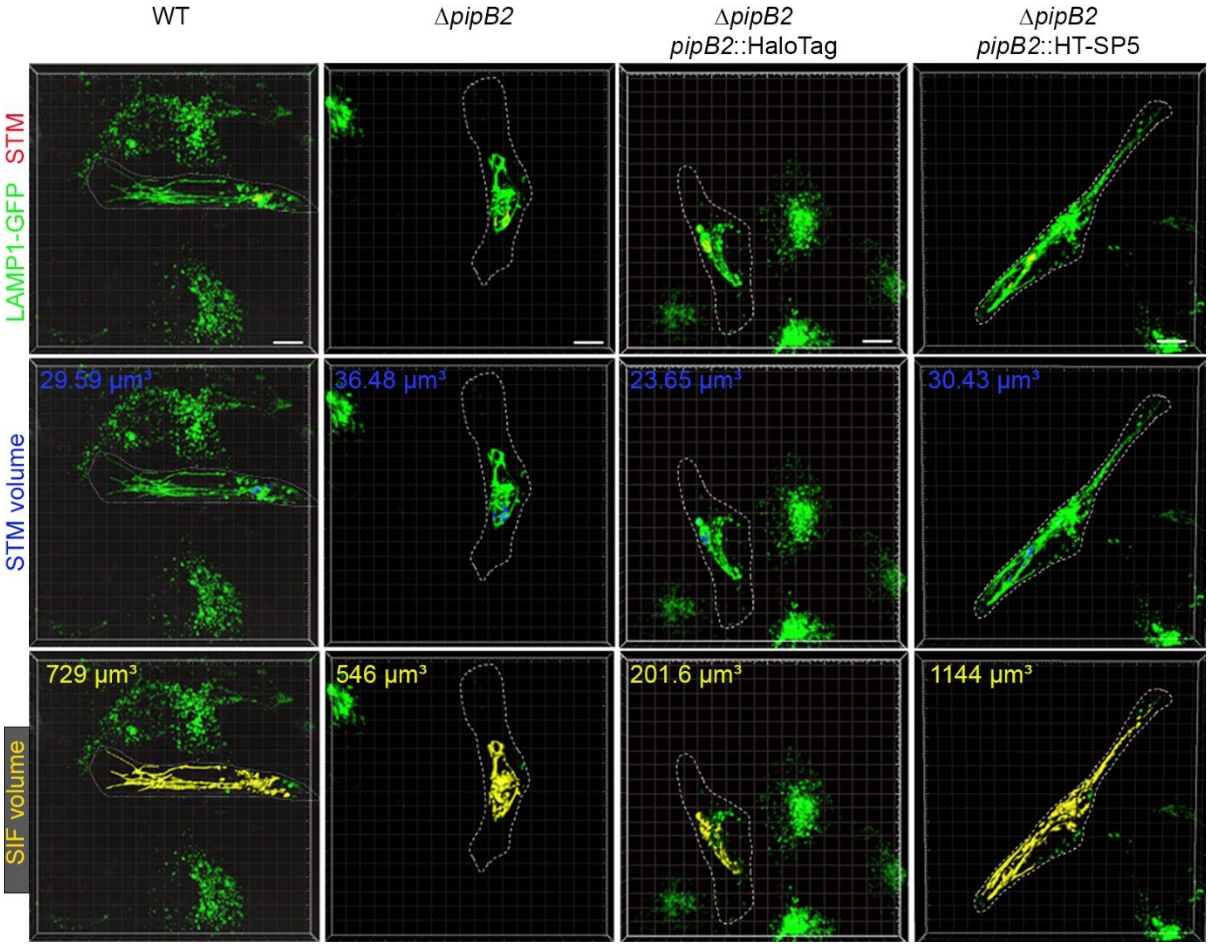


**Figure S1: Effect of distinct point mutations in HaloTag on the translocation of functional effector-HaloTag fusions involved in SIF formation.** HeLa LAMP1-GFP cells (green) were infected with the STM WT or the STM Δ*pipB2* strain expressing sfmTurquoise2ox. Plasmids for expression of *pipB2*::HaloTag::HA or *pipB2*::HT-SP5::HA were used for complementation. 3D reconstructions of representative cells and segmentations of volumes of LAMP1-GFP-positive SCV enclosing *Salmonella* (blue) and interconnected SIF tubules (yellow) Scale bars, 10 µm.

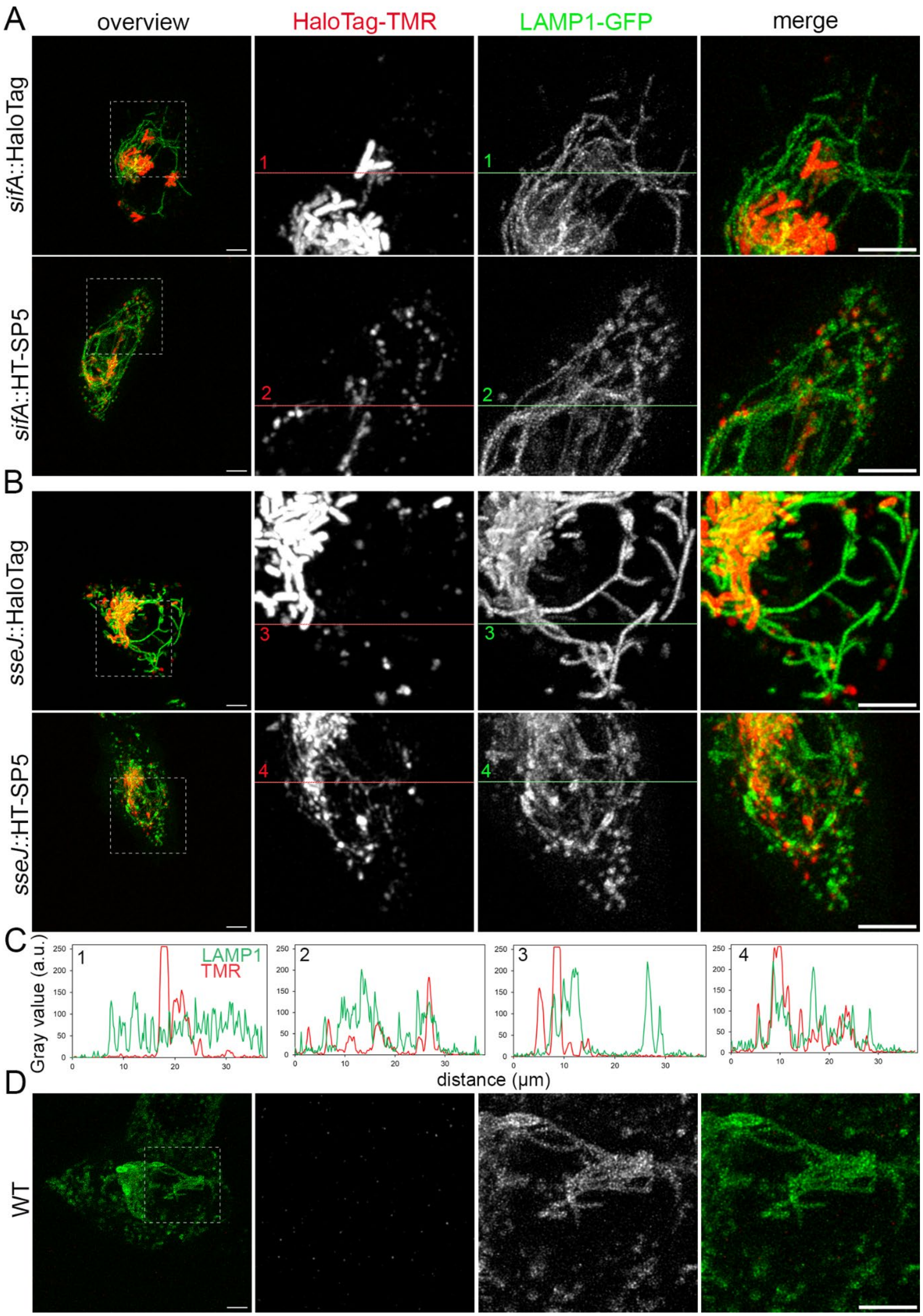


287

288 **Figure S2: Enhanced translocation of SPI2-T3SS effector proteins fused to HT-SP5**. For
289 analyses of effector translocation, HeLa cells stably expressing LAMP1-GFP were infected by

STM WT harbouring plasmids for the expression of SifA **(A)** or SseJ **(B)** fused to HaloTag or HT-SP5. STM WT with empty vector served as negative control (**D)**. At 8 h p.i., labelling with 1 µM HTL-TMR was performed for 30 min at 37 °C directly before imaging. Representative STM-infected host cells exhibiting SIF formation were selected for LCI. Scale bars, 10 µm. (**C**) Line scans reveal intra-bacterial and translocated effector quantities in relation to LAMP1-GFP signal intensities.

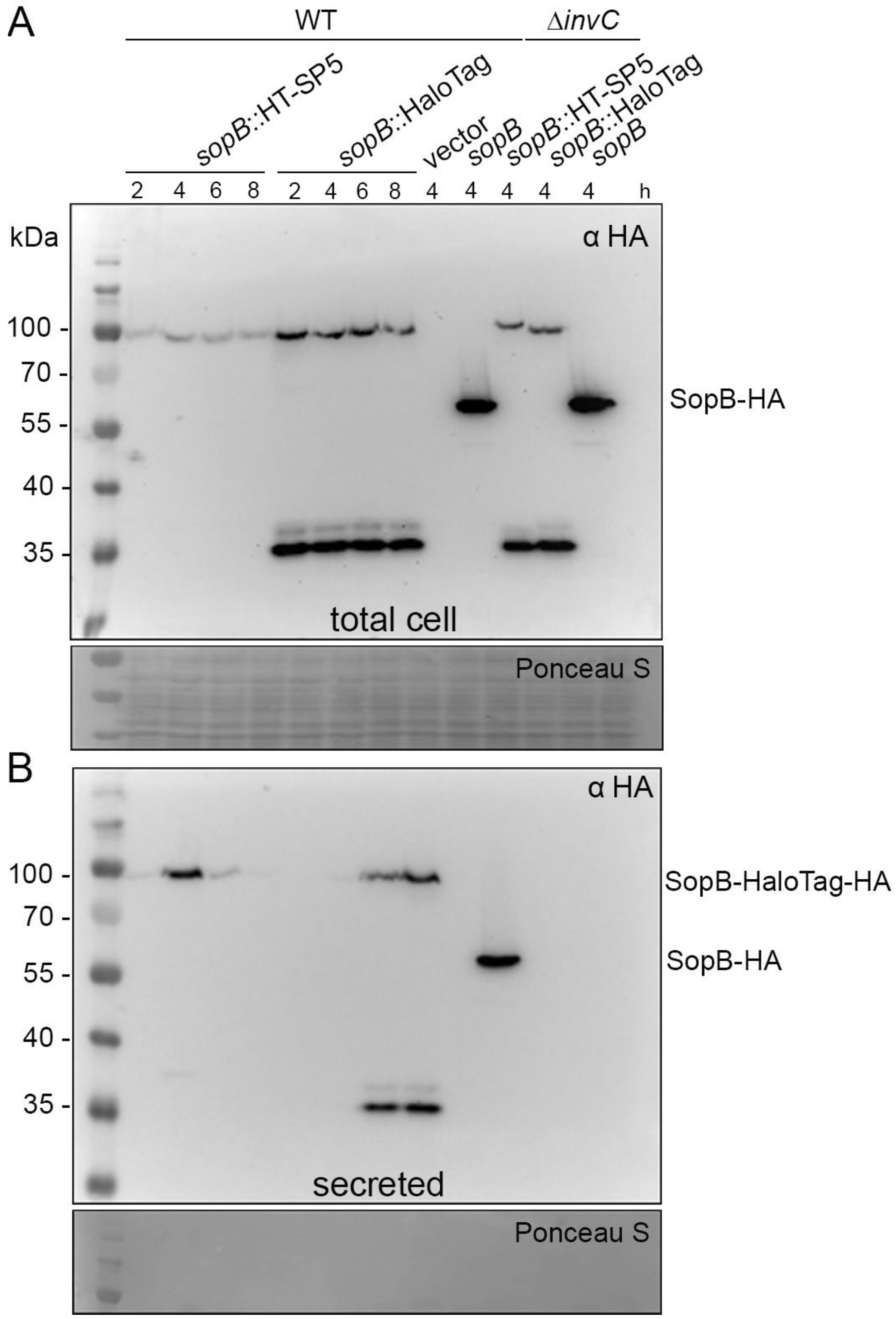


**Figure S3: Synthesis and secretion of SopB-HaloTag fusions.** STM WT or SPI1-T3SS-deficient strain Δ*invC*, harbouring empty vector or plasmids for expression of *sopB*::HA, *sopB*::HaloTag::HA or *sopB*::HT-SP5::HA, were grown with aeration at 37 °C in LB. At indicated time points, bacterial pellets **(A)** and supernatants **(B)** were collected and analysed by Western blot using anti-HA antibodies. The Ponceau S-red staining is shown below as loading control. Detected bands represent SopB-HA (63.4 kDa), SopB-HaloTag-HA (96.8 kDa).

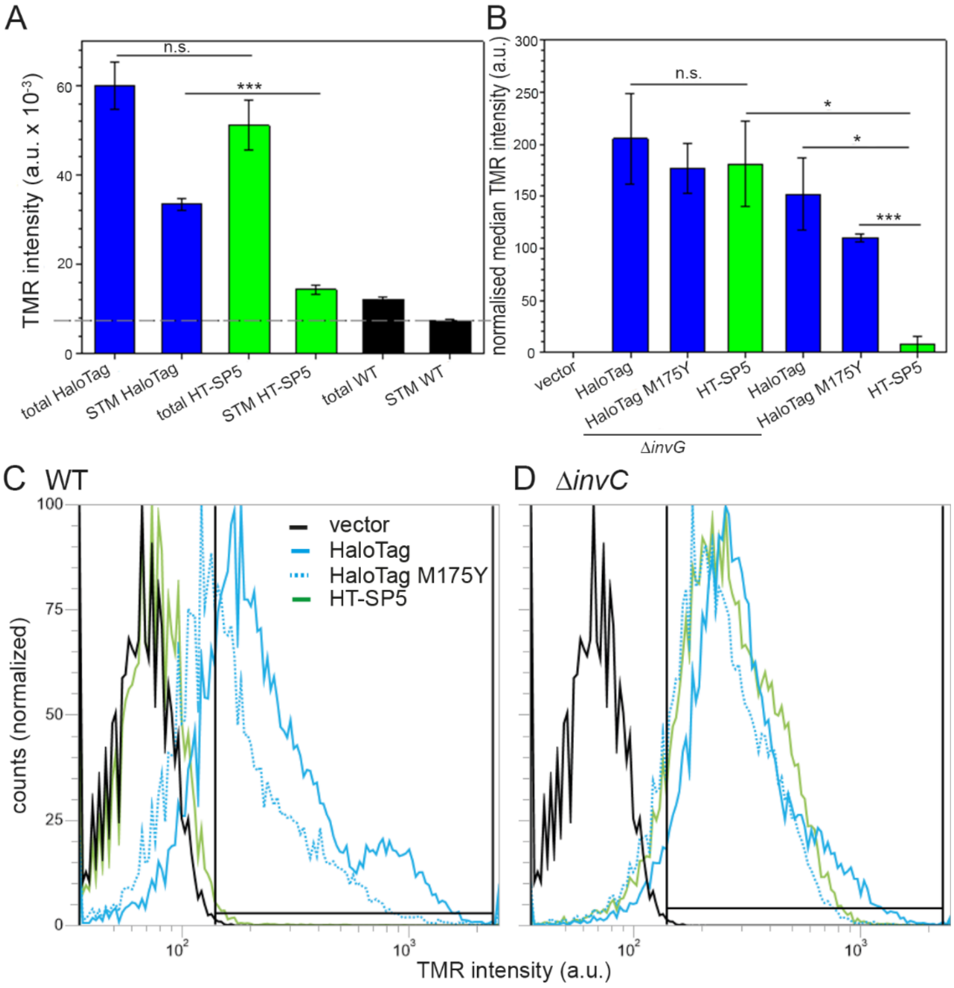


**Figure S4: Alleles of HaloTag show comparable fluorescence intensities upon HTL-TMR labelling but differential effector translocation and secretion potential. A)** RAW macrophages were seeded in 6-well plates and infected with STM WT expressing *pipB2*::HaloTag and *pipB2*::HT-SP5 at MOI 5. 8 h p.i., 1 µM HTL-TMR was added for 30 min before cells were detached or lysed. 200 µl of cells and lysate were measured using a microplate reader (Infinite 200 Pro M-Plex, Tecan). **B-D**) STM WT and a secretion-deficient mutant (Δ*invG*) without or with plasmids for expression of *sopE*::HaloTag fusions were grown for 3.5 h in LB medium. Equal bacterial amounts were labelled with HTL-TMR for 40 min at 37 °C. After washing to remove unbound ligand and secreted effectors, at least 25,000 bacteria-sized particles were analysed by flow cytometry, and TMR fluorescence was determined as proxy for cell-bound, non-secreted amounts of SopE. Normalised median fluorescence intensity of the gated population was calculated from three biological replicates. **C, D**) The gating strategy and TMR intensity distribution.

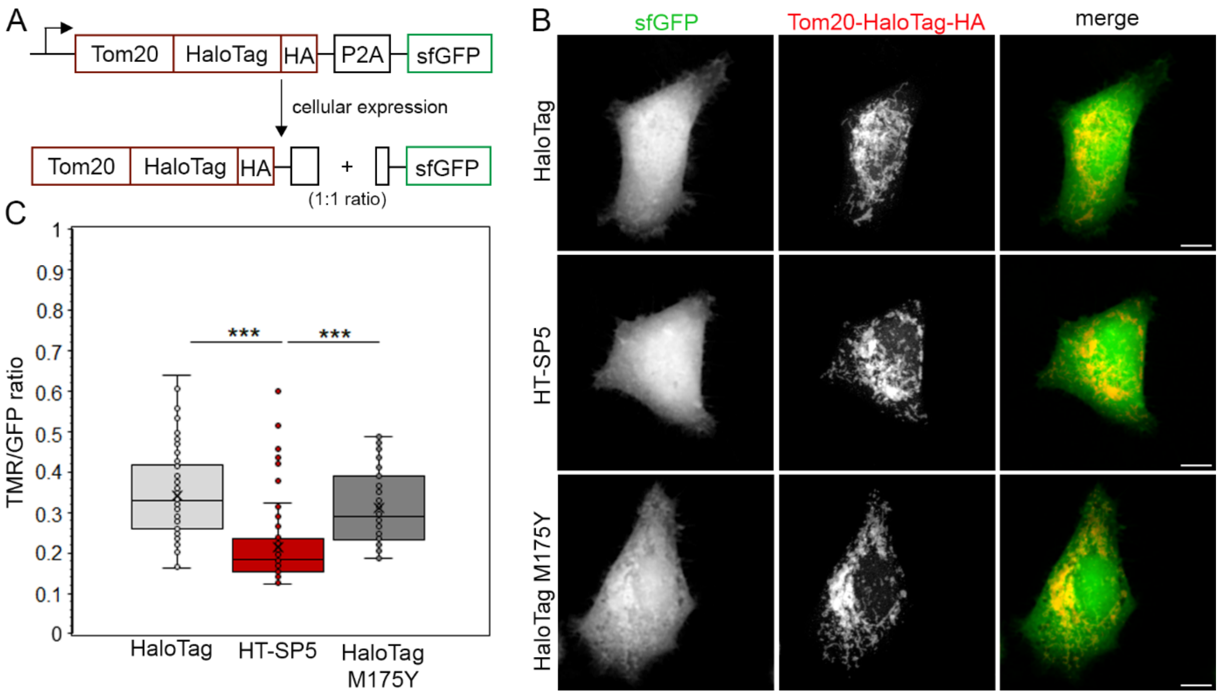


**Figure S5: Effect of the various HaloTag fusions on the labelling efficiency by HTL-TMR. A, B)** HeLa cells were transiently transfected for expression of Tom20::HaloTag::HA::P2A::sfGFP for the indicated HaloTag alleles. At 19 h after transfection, cells were labelled with 100 nM HTL-TMR for 30 min at 37 °C, fixed, and imaged by cLSM. Scale bars, 10 µm. **C)** Quantification of TMR and sfGFP signals was performed using ImageJ 1.53t. Statistical significances were determined using an unpaired, two-tailed Student's t-test and indicated as: ***, $p < 0.001$. Box plots represent data from 72 analysed cells per HaloTag variant, with X and horizontal lines within boxes representing means and medians, respectively, of calculated TMR/GFP ratios.

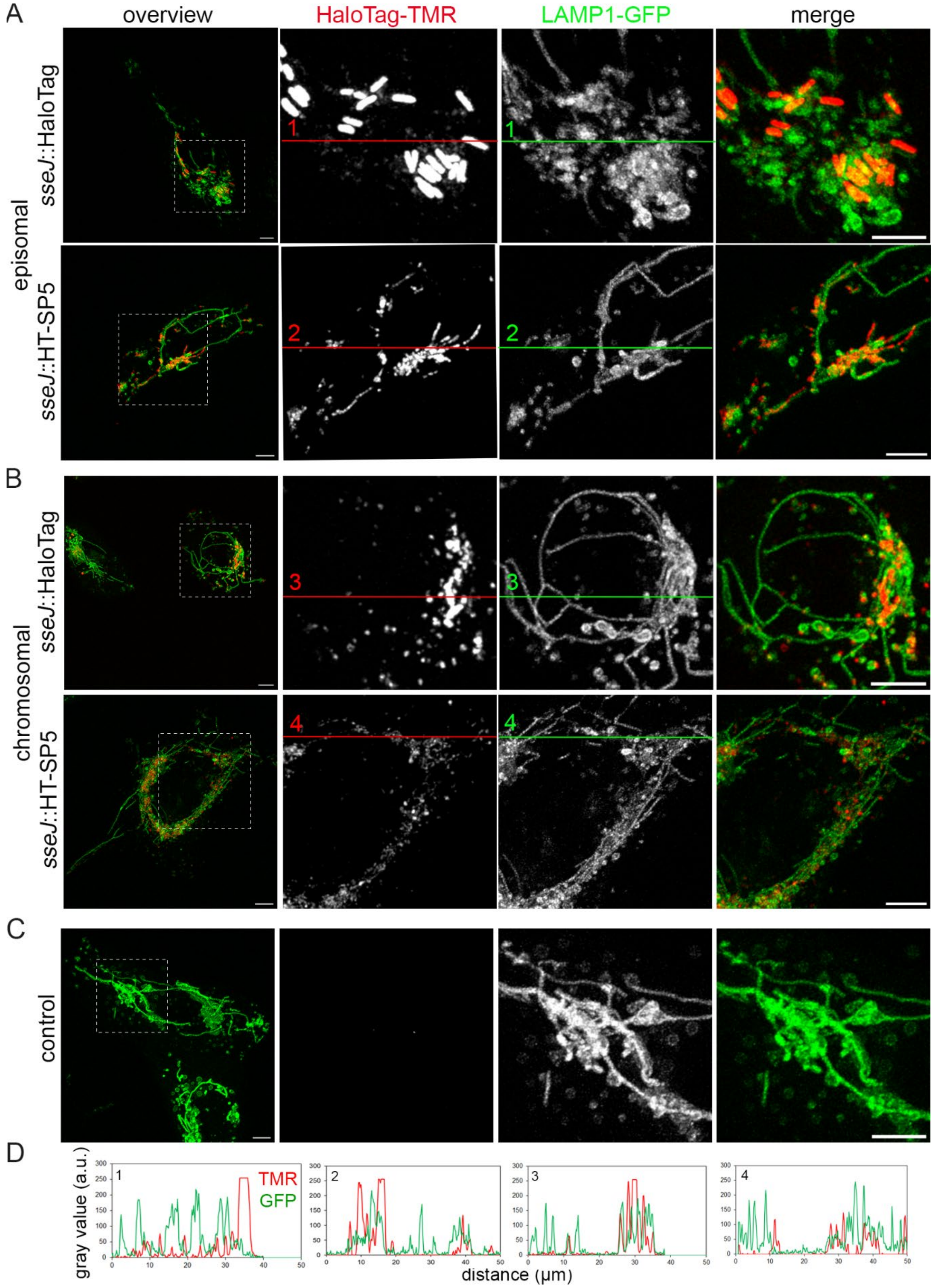


**Figure S6: Efficient translocation of chromosomal encoded effector-HT-SP5 fusions.** HeLa cells stably expressing LAMP1-GFP were infected with STM strains expressing lowcopy plasmid-encoded **(A)** and chromosomal **(B)** fusions of *sseJ* to HaloTag or HT-SP5 at MOI 10. After labelling with 1 µM HTL-TMR (red) for 30 min at 37 °C, LCI was performed. STM WT with empty vector served as negative control **(C)**. Scale bars, 10 µm. **D)** Line scans show the translocated and intra-bacterial SseJ signal in relation to the LAMP1-GFP signal.

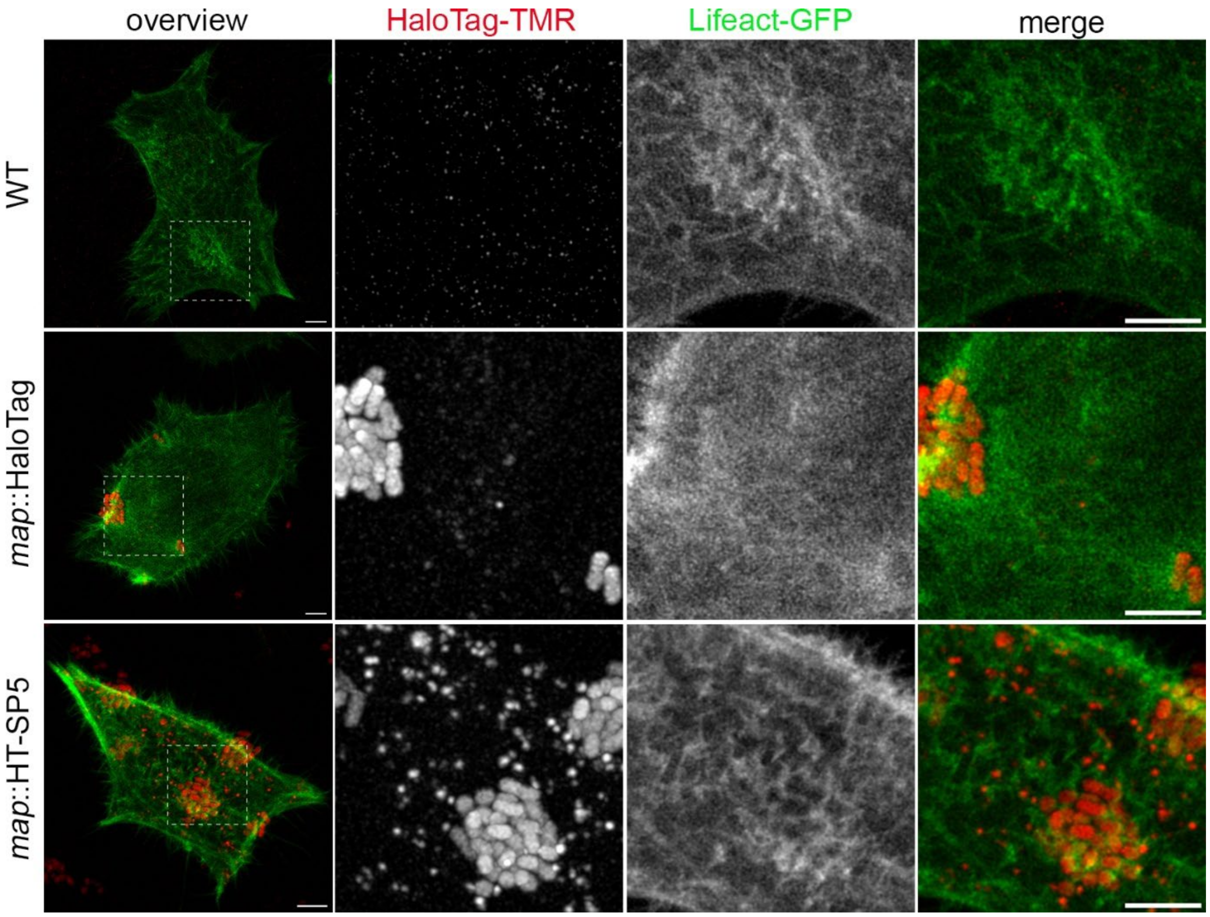


**Figure S7: Enhanced translocation of enteropathogenic *E. coli* (EPEC) T3SS effector protein Map fused to HT-SP5.** For translocation of EPEC effector protein Map, infection of HeLa cells constitutively expressing Lifeact-GFP (green) was performed with EPEC E2348/69 harbouring plasmids for the expression of *map* fused to HaloTag or HT-SP5 as indicated. EPEC WT was used as a negative control. At two h p.i., labelling reactions were performed directly before imaging using a final concentration of 0.8 µM HTL-TMR for 30 min at 37 °C. Scale bars, 10 µm.

### Movie captions

**Movie S1: Translocation of SopE-HT-SP5.** HeLa cells stably expressing Lifeact-GFP were infected with STM WT expressing *sopE*::HT-SP5 and mNeonGreen. After centrifugation for 5 min to synchronize infection, time-lapse microscopy using a LLSM was performed in 60 sec intervals in the presence of 20 nM MaP555. 3D projections of image stacks were generated (Imaris) and annotated (Adobe Premiere Pro). Note the appearance of MaP555-positive spots in host cells at sites of invading STM.

**Movie S2: Translocation and localization dynamics of PipB2-HT-SP5.** HeLa cells stably expressing LAMP1-GFP were infected with STM WT expressing AHT-inducible *pipB2*::HT-SP5 and mNeonGreen. Time-lapse imaging in the presence of 20 nM MaP555 and 50 nM AHT was performed using LLSM starting 4 h p.i. within 60 sec intervals. 3D projections of image stacks were generated (Imaris) and annotated (Adobe Premiere Pro). Note the appearance of MaP555-positive spots in host cells at sides of intracellular STM and the integration of vesicles double-positive for LAMP1 and MaP555 into the dynamic SIF network.

**Movie S3: Controls for translocation of effector fusions to HT-SP5.** HeLa cells stably expressing Lifeact-GFP remain uninfected **(A)** or were infected with STM WT expressing mNeonGreen (**B**). HeLa cells stably expressing LAMP1-GFP were infected with STM Δ*ssaV* expressing mNeonGreen and PipB2-HT-SP5 (**C**) and STM WT expressing mNeonGreen and PipB2-HT-SP5 (**D**). After centrifugation for 5 min to synchronize infection (**B**), or 4 h p.i. (**C, D**), time-lapse imaging using a LLSM system in presence of 20 nM MaP555 was performed every 60 sec. **C**) 4 h p.i., 50 ng/ml AHT was added to induce expression of *pipB2*::HT-SP5. 3D projections of image stacks were generated (Imaris) and annotated (Adobe Premiere Pro). Note the absence of MaP555-positive spots in host cells in the absence of translocated effector-HT-SP5-fusions.
